# Stress-hardening behaviour of biofilm streamers

**DOI:** 10.1101/2024.10.04.616620

**Authors:** Giovanni Savorana, Tommaso Redaelli, Domenico Truzzolillo, Luca Cipelletti, Eleonora Secchi

## Abstract

Bacteria’s ability to withstand mechanical challenges is enhanced in their biofilm lifestyle, where they are encased in a viscoelastic polymer matrix [1]. Under fluid flow, biofilms can form as streamers – slender filaments tethered to solid surfaces and suspended in the flowing fluid [2, 3]. Streamers thrive in environments subjected to intense hydrodynamic stresses, such as medical devices and water filters, often resulting in catastrophic clogging [4]. Their colonisation success may depend on a highly adaptable mechanical response to varying stress conditions, though the evidence and underlying mechanisms of this adaptation remain elusive. Here, we demonstrate that biofilm streamers exhibit a stress-hardening behaviour, with both differential elastic modulus and effective viscosity increasing linearly with external stress. This stress-hardening is consistent across biofilms with different matrix compositions, formed by various bacterial species, and under diverse growth conditions. We further demonstrate that this mechanical response originates from the properties of extracellular DNA (eDNA) molecules [5], which constitute the structural backbone of the streamers. In addition, our results identify extracellular RNA (eRNA) as a modulator of the matrix network, contributing to both the structure and rheological properties of the eDNA backbone. Our findings reveal an instantaneous, purely physical mechanism enabling streamers to adapt to hydrodynamic stresses. Given the ubiquity of extracellular nucleic acids (eNA) in biofilms [1, 6], this discovery prompts a re-evaluation of their functional role in biofilm mechanics, with potential implications for biofilm structural integrity, ecological resilience, and colonisation dynamics.

## Introduction

Biofilms are ubiquitous bacterial communities held together by a matrix of extracellular polymeric substances (EPS) [1, 7, 8]. Their ability to thrive in diverse environments is largely attributed to the EPS matrix’s viscoelastic behaviour [9–13], which is also recognised as a virulence factor in biofilm-associated infections [10]. Protected by the matrix, the biofilm community adapts to mechanical stresses from the environment through both biological and physical processes. On the one hand, mechanical stresses can activate mechanosensing responses at the single-cell level, which in turn regulate EPS secretion [14, 15]. On the other hand, physical interactions between bacteria and their surroundings can induce cellular ordering and shape the biofilm microstructure [16–18]. Additionally, preliminary studies suggest that biofilms may exhibit nonlinear rheological behaviours under stress [19–23]. However, it remains unclear whether and how biofilms can adjust their mechanical properties in response to environmental mechanical stresses.

Biofilms can successfully colonise environments ranging from medical devices to natural ecosystems, where flow velocities and hydrodynamic stresses can vary by several orders of magnitude [11]. Notably, fluid flow promotes the formation of streamers, slender biofilm filaments tethered to surfaces and suspended in the bulk of the flow [4, 24]. The formation and structural integrity of streamers rely entirely on the viscoelastic nature of the EPS matrix, which enables them to withstand hydrodynamic stresses and extend far from colonised surfaces, facilitating bacterial spreading [2, 3]. Streamer formation has been mainly studied in *Pseudomonas aeruginosa*, a key pathogen in chronic infections and antibiotic resistance, and a model organism for biofilm research [25–29]. In the PA14 strains, the main EPS components of streamers are extracellular DNA (eDNA), which constitutes their structural backbone, and the polysaccharide Pel, which influences their morphology and viscoelastic properties [2, 3]. eDNA provides structural integrity also to *P. aeruginosa* PAO1 and *Burkholderia cenocepacia* streamers [3, 30]. Evidence for its structural role includes the DNase I–induced disintegration of streamers in *P. aeruginosa* PA14 [3], and the lack of streamer formation in *B. cenocepacia* mutants lacking the genetic pathway that mediates eDNA release via cell lysis [30]. Given that single DNA molecules exhibit stress-hardening behaviour – thus stiffen in response to increasing mechanical stress [5, 31] – it can be hypothesised that eDNA not only contributes to the structural integrity of streamers but also enhances their mechanical resilience.

DNA is not only present in streamers, but it is also recognized as a key structural element in surface attached single- and multi-species biofilms [6, 32–35]. In these systems, its highly charged nature enables interactions with other matrix components. Such interactions include binding with polysaccharides [3, 27, 36], and with DNA-binding proteins of the DNABII family [37, 38], which are increasingly recognized as critical for biofilm stability and as promising targets for biofilm eradication [39]. While eDNA has been widely acknowledged as a critical matrix component, recent studies also implicate extracellular RNA (eRNA) in biofilm structuring [40, 41]. eRNA appears to stabilise eDNA fibres in *P. aeruginosa* biofilms and influence their viscoelastic properties by promoting the formation of eDNA supramolecular structures such as Holliday junctions [41]. However, most of this evidence comes from studies on surface-attached biofilms and relies on pretreated or isolated nucleic acid networks, while the structural role of eDNA and eRNA remains largely unexplored *in situ* and within the eDNA-rich suspended biofilm streamers.

In this work, we characterise biofilm streamers grown under a range of flow velocities to investigate how their viscoelastic properties change in response to the hydrodynamic stress exerted by the environment. We identify and model the primary mechanism driving their mechanical adaptation, which we found to be conserved across multiple bacterial species, and determined by the extra-cellular nucleic acids (eNA) in the matrix. Identifying biofilm adaptive mechanical response and uncovering its underlying mechanism advances our understanding of biofilm resilience in dynamic environments, offering novel perspectives on biofilm ecology and providing crucial insights for the development of more effective eradication strategies.

### *In situ* characterisation of biofilm streamer viscoelasticity

To systematically examine the viscoelasticity of biofilm streamers under different flow velocities, we use a microfluidic platform to grow streamers with a reproducible morphology and characterise their material properties *in situ* [2]. Pillar-shaped obstacles placed in a straight microfluidic channel act as nucleation points for pairs of biofilm streamers, which develop as millimetre-long filaments tethered to the sides of the pillars when a diluted bacterial suspension is flowed through the channel. They grow over time, extending in the flow direction. Once the streamers reach a steady state, we stain them with propidium iodide (PI) and image them using epifluorescence microscopy. PI binds to nucleic acids, primarily DNA, and enables the reconstruction of the streamers’ three-dimensional geometry. The reconstruction serves as the basis for computational fluid dynamics (CFD) simulations, which allow us to estimate the forces exerted by the flow on the streamers [2]. Then, we perform extensional rheological measurements on nearly cell-free portions of streamers, thus determining the viscoelastic properties of homogeneous EPS matrix samples. Since streamers develop under continuous background flow, they are constantly subjected to an extensional axial stress *σ* and remain in a state of deformation with a non-zero extensional strain *ε*. The axial stress at position *x* along a streamer of total length *L* depends on the surface area of the downstream portion of the streamer exposed to flow and is expressed as:

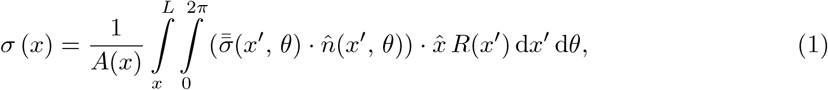

where *A* (*x*) = *πR*^2^ (*x*) is the cross-sectional area of the streamer at *x*, 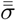 is the Cauchy stress tensor of the fluid flowing past the streamer, 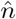 is the unit vector perpendicular to the streamer surface, and 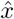 is the unit vector in the *x* direction, parallel to the main axis of the streamer [2]. Equation 1 shows that the axial stress *σ*(*x*), at a given position *x* along the streamer, depends not only on the stress tensor of the fluid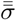, which is proportional to the flow velocity, but also on the morphology of the portion of streamer downstream of *x*. To test the viscoelastic properties of a streamer, we apply a controlled flow perturbation that imposes an increase in extensional stress Δ*σ* on top of the prestress Δ*σ*_0_ corresponding to the unperturbed flow velocity. This results in an additional filament deformation characterised by a strain increment Δ*ε*. From these increments, we calculate the differential Young’s modulus *E*_diff_ and effective viscosity *η*, quantifying the material properties of the streamer as a function of its prestress state *σ*_0_. This differential testing method is analogous to the one implemented for characterising the shear rheology of cytoskeletal and extracellular biological networks in tissues [42, 43].

### Ambient flow affects biofilm streamer morphology

We investigate the adaptation of streamer morphology, biochemical composition, and viscoelasticity to different flow velocities *U*_gr_ using *P. aeruginosa* PA14. To investigate the role of polysaccharides in the adaptation process, we compare PA14 strains with different levels of Pel polysaccharide production: the wild-type (wt) strain, which produces Pel in response to external forces [14, 15], the Pel-deficient mutant (Δpel) and the Pel-overproducer (ΔwspF), which is locked in the active state of Pel production [3].

Microfluidic experiments reveal that all *P. aeruginosa* PA14 strains form biofilm streamers across a range of flow velocities spanning an order of magnitude within the laminar regime (Re ∈ [0.02, 0.20], Supplementary Table 1). After 15 hours, streamers formed under different velocities exhibit similar morphologies but vary in average length *L* and radius *R* (Fig. 1e-f). These differences are due to flow-induced changes in cell and EPS assembly rather than biomass flux, which we maintain constant across conditions. The average length notably decreases with increasing flow velocity, regardless of Pel production levels (Fig. 1e). Time-lapse imaging shows that higher velocities delay the formation of the first stable biofilm threads, likely due to reduced cell attachment, consistent with observations on flat surfaces [44] (Supplementary Figure 1a, b). After the initial time lag *t*_0_, the length increases with time approximately following a power law behaviour, *L ∝* (*t − t*_0_)^0.5^, under all flow conditions, with slower growth rates at higher velocities (Supplementary Figure 1c). Near the pillar (ROI_1_, Fig. 1a, c), the radius increases significantly with flow velocity (Fig. 1f), while downstream (ROI_2_, Fig. 1b, d), streamers are typically thinner, with the radius increasing with flow velocity for the wild-type strain, but showing no clear trends for the Pel-deficient and overproducer strains (Fig. 1f). Overall, these findings indicate that flow velocity significantly influences streamer development and morphology, with limited dependence on Pel polysaccharide abundance. It is important to highlight that the dependence of streamer morphology on ambient flow conditions implies that streamers grown at higher flow velocities are not necessarily subjected to higher extensional axial stresses, according to eq. 1.

**Fig. 1.**
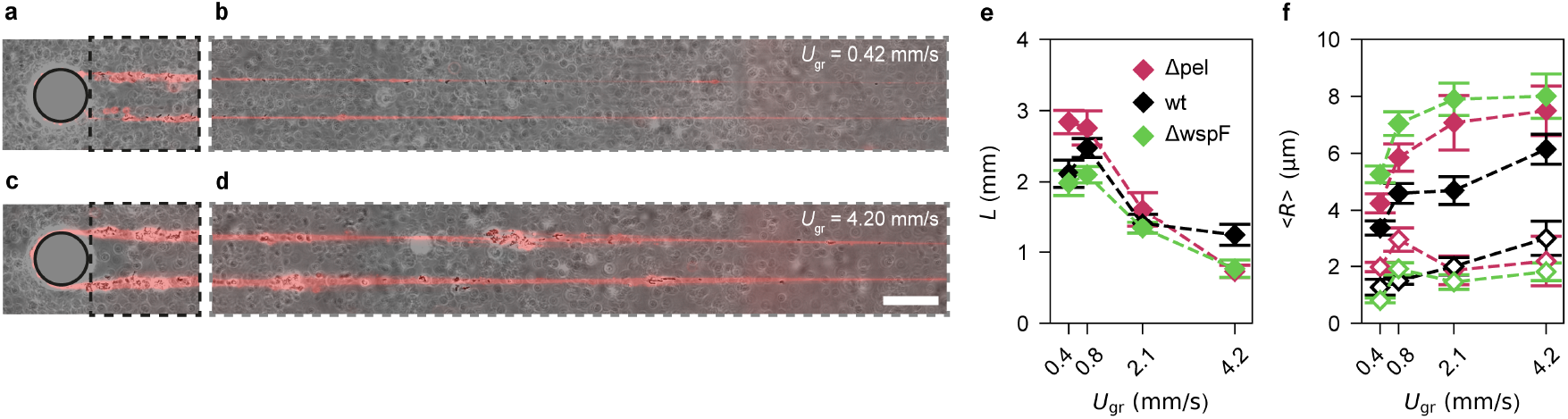
The morphology of *P. aeruginosa* PA14 biofilm streamers is affected by ambient flow conditions during their growth. **a**-**d**, Representative overlaid phase contrast and fluorescence images of 15-hour-old PA14 wild-type biofilm streamers grown on a micropillar at the lowest (*U*_gr_ = 0.42 mm*/*s, **a, b**), and highest (*U*_gr_ = 4.20 mm*/*s, **c, d**) average flow velocities tested in this study. The flow is laminar, and the flux of bacterial cells is constant across the different flow conditions. The black and grey dashed rectangles highlight the regions corresponding to ROI_1_ (*x* ∈ [25 µm, 125 µm]) and ROI_2_ (*x* ∈ [400 µm, 1065 µm]), respectively. The scale bar is 50 µm. Bacterial cells were imaged in phase contrast, and nucleic acids were stained using red-fluorescent propidium iodide (2 µg*/*mL). **e**-**f**, Average length *L* (**e**) and average of ⟨*R*⟩ in ROI_1_ (**f**, closed markers) and ROI_2_ (**f**, open markers) as a function of the average flow velocity *U*_gr_ inside the microchannel during streamer formation for PA14 Δpel (magenta), wild-type (black) and ΔwspF (green). The error bars are the standard error of the mean.

### Rheological characterisation of streamers grown under different flow velocities

Mechanical tests performed on PA14 biofilm streamers grown under different flow velocities show a linear relationship between the matrix viscoelastic properties and the axial stress exerted by fluid flow, revealing a strong mechanical adaptation mechanism of streamer viscoelasticity. We demonstrate this by growing biofilm streamers for 15 hours at a constant flow velocity, and then characterising the rheology of specific portions delimited by pairs of cell aggregates located at positions *x*_1_ and *x*_2_ within ROI_2_. During the mechanical tests, we instantaneously double the initial flow velocity from *U*_0_ = *U*_gr_ to *U*_cr_ = 2*U*_0_ (Supplementary Figure 2a). This increase results in a change in the average axial stress on the examined portion of the streamer from the prestress value *σ*_0_ ≡ ⟨*σ*(*x*; *U*_0_)⟩_12_ to the creep value *σ*_cr_ ≡ ⟨*σ*(*x*; *U*_cr_)⟩_12_, where the average is calculated between *x*_1_ and *x*_2_, and the notation *σ*(*x*; *U*) explicitly shows the parametric dependence of the stress on flow velocity *U*. Neglecting fluid-structure interaction [2], we have *σ*_cr_ = 2*σ*_0_ (Supplementary Figure 2b). The corresponding stress increment Δ*σ* = *σ*_cr_ −*σ*_0_ is maintained for 5 minutes, during which we use optical microscopy to quantify the time-dependent strain increment Δ*ε* (*t*), consisting of an instantaneous elastic contribution Δ*ε*_el_ and an irreversible time-dependent viscous deformation with constant strain rate ε̇ = d (Δ*ε* (*t*)) */*d*t* (Supplementary Figure 2c). Finally, we use Δ*σ* and Δ*ε* (*t*) to determine the differential Young’s modulus *E*_diff_ = Δ*σ/*Δ*ε*_el_ and the effective viscosity *η* = Δσ/ε̇ of the sample portion under the prestress condition *σ*_0_. Our experiments reveal that increasing the prestress exerted by ambient flow leads to enhanced mechanical resistance of the streamers, as demonstrated by a marked increase in differential Young’s modulus and effective viscosity with prestress. Across the wide range of conditions tested (Supplementary Figure 3), both quantities exhibit an approximately linear correlation with prestress (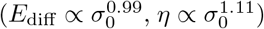) (Fig.2).

We find that the observed linear dependence of the differential Young’s modulus and effective viscosity on the prestress is independent of the EPS matrix composition in PA14 biofilm streamers. First, this trend is not determined by variations in eDNA density, which remains constant across a wide range of prestress conditions (Supplementary Figure 4). Second, we observe the same linear dependence of differential Young’s modulus and the effective viscosity on prestress in the PA14 wt, Pel-deficient, and Pel-overproducer strains. This demonstrates that the presence or abundance of Pel does not determine whether stress-hardening occurs, nor the functional dependence of viscoelastic properties on prestress. However, Pel abundance modulates the magnitude of the viscoelastic properties at a given prestress (Supplementary Fig. 5), in agreement with our previous work [2, 3]. Specifically, for a given prestress value, the Pel-overproducer strain consistently displays a higher differential Young’s modulus than both the Pel-deficient and the wild-type strains, which show similar values (Supplementary Figure 5b, c). Pel staining reveals that in the wild-type strain, significant amounts of Pel are detected only near the pillar (ROI_1_ in Supplementary Fig. 6a-d) at higher flow velocities (*U*_gr_ = 2.10 ml*/*h and *U*_gr_ = 4.20 ml*/*h), while Pel concentration in the region where the mechanical properties are probed (ROI_2_ in Supplementary Figure 6a-d) remains below our detection limit, regardless of flow velocity. Given that the viscoelastic properties of the EPS matrix correlate with the spatial distribution of polysaccharides within the biofilm [45], the similarity in the viscoelastic properties measured in both the wild-type and Pel-deficient mutants of the biofilm is consistent with the low levels of Pel polysaccharide found in the wild-type strain. In contrast, the ΔwspF mutant shows detectable Pel concentration in both ROI_1_ and ROI_2_ across all flow conditions, explaining its higher differential Young’s modulus and effective viscosity compared to the other strains at a given prestress (Supplementary Figure 6e-h).

Given that the linear relationship between the viscoelastic properties and the prestress measured on streamers grown under different flow velocities cannot be attributed to flow-induced differences in EPS matrix composition, which are observed in surface-attached biofilms [17], we hypothesise that it may result from a nonlinear response of the matrix to mechanical stresses, as previously observed in eukaryotic biological networks formed from cytoskeletal and extracellular proteins [42, 43, 46]. To validate this hypothesis, each streamer must be tested under multiple prestress conditions.

### Stress-hardening behaviour in PA14 biofilm streamers

By testing the rheology of each streamer under multiple prestress conditions *σ*_0_, we demonstrate that the linear dependence of the viscoelastic properties on prestress results from the stress-hardening behaviour of the EPS matrix. To conduct the tests, we first grow PA14 streamers at *U*_gr_ = 2.10 mm*/*s for 15 h. After this growth period, we perform two sequential mechanical tests on each sample. The tests are conducted using different initial flow velocities *U*_0_ for each test, with each *U*_0_ having a corresponding creep velocity *U*_cr_ = 2*U*_0_. During the first test, the initial flow velocity is set to *U*_0_ = *U*_gr_, which corresponds to a prestress of *σ*_0_ = ⟨*σ*(*x*; *U*_gr_)⟩_12_, as in the tests presented in the previous paragraph. After the first test, we decrease the flow velocity to *U*_gr_*/*5 = 0.42 mm*/*s and let the system stabilise for 10 minutes. Following this, we perform a second mechanical test with initial flow velocity *U*_0_ = *U*_gr_*/*5, corresponding to a prestress *σ*_0_ = ⟨*σ*(*x*; *U*_gr_*/*5)⟩_12_ (Supplementary Figure 2). We image and analyse the same portion of the streamer in both tests to measure the differential Young’s modulus and effective viscosity of the same sample under the two different prestress conditions (Supplementary Figure 2d-f). To ensure that the observed viscoelastic properties are independent of the loading history, in selected experiments, we reverse the order of the mechanical tests and probe the streamer first under prestress *σ*_0_ = ⟨*σ*(*x*; *U*_gr_*/*5)⟩_12_ and then under prestress *σ*_0_ = ⟨*σ*(*x*; *U*_gr_)⟩_12_.

The differential Young’s modulus and effective viscosity values measured sequentially on the same sample exhibit the same linear dependence on prestress as in streamers grown under different ambient flow conditions (Fig. 2, 3). This trend is consistent across all PA14 mutants, regardless of their Pel polysaccharide production levels, and is, therefore, independent of Pel abundance. The 10-minute interval between the tests is sufficient for the system to stabilise at the new flow velocity but too brief for bacteria to initiate any adaptive mechanisms [14]. Therefore, we conclude that the rheological adaptation of the streamer EPS matrix to the prestress is an instantaneous, purely physical stress-hardening response, which also explains the trends observed for streamers grown under different flow velocities (Fig. 2).

**Fig. 2.**
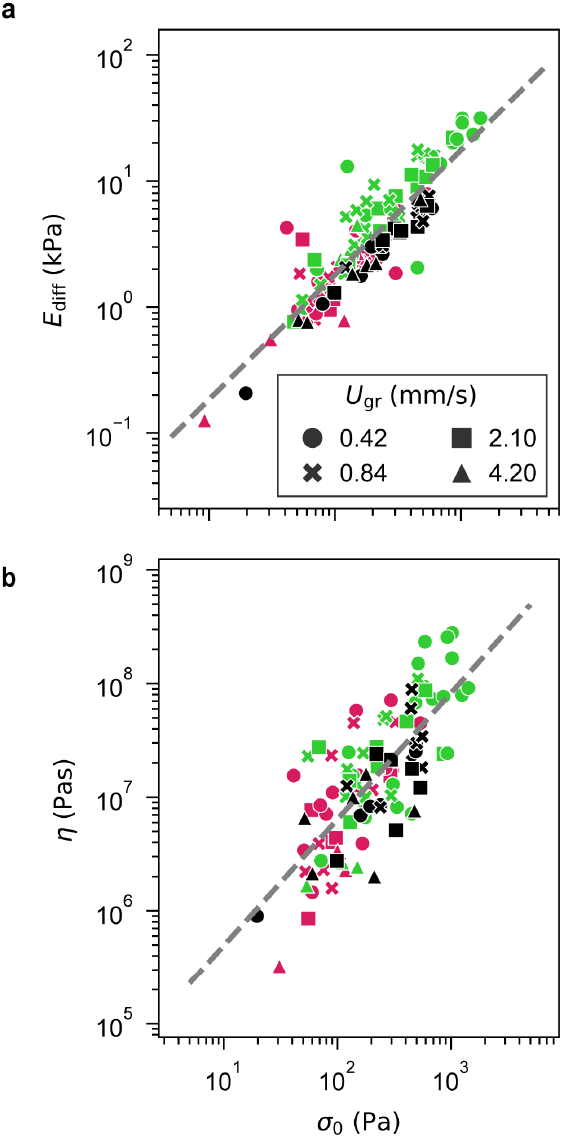
The viscoelastic properties of *P. aeruginosa* PA14 biofilm streamers linearly depend on the prestess *σ*_0_ **exerted by** *U*_gr_. **a, b**, Differential Young’s modulus *E*_diff_ and effective viscosity *η* as a function of prestress *σ*_0_. Each point is measured on a portion of a different streamer, whose prestress value *σ*_0_ = ⟨*σ*(*x*; *U*_gr_)⟩_12_ is determined by sample morphology and *U*_gr_. All samples are tested after 15 hours of growth. Different colours represent the different PA14 mutants: Δpel (magenta), wild-type (black), and ΔwspF (green). Different symbols represent data points obtained for samples grown at different average flow velocities *U*_gr_: 0.42 mm*/*s (circles), 0.84 mm*/*s (crosses), 2.10 mm*/*s (squares) and 4.20 mm*/*s (triangles). The dashed grey lines are power-law fits of the data, with exponent 0.99 *±* 0.05 (*R*^2^ = 0.79) with a 95% confidence interval [0.90, 1.08] in **a** and 1.11 *±* 0.10 (*R*^2^ = 0.56) with a 95% confidence interval [0.92, 1.31] in **b**.

**Fig. 3.**
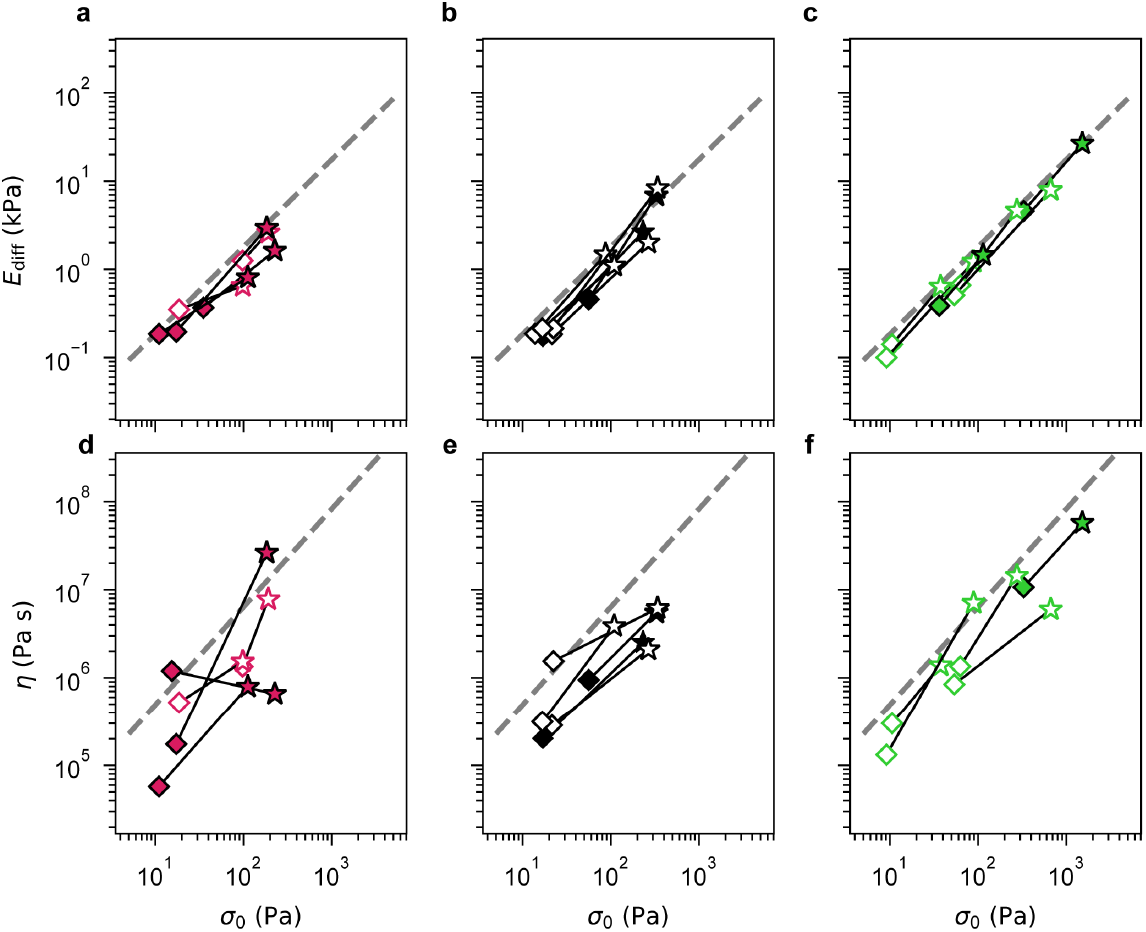
The EPS matrix of *P. aeruginosa* PA14 biofilm streamers has stress-hardening viscoelastic properties. **a**-**f**, Differential Young’s moduli *E*_diff_ (**a**-**c**) and effective viscosities *η* (**d**-**f**) obtained from sequential mechanical tests demonstrate the stress-hardening behaviour of the matrix of PA14 Δpel (**a, d**, magenta), wild-type (**b, e**, black), and ΔwspF (**c, f**, green) streamers. Each pair of markers connected by a solid line represents values sequentially measured on a same sample under two different prestress conditions *σ*_0_. The tests are conducted on PA14 biofilm streamers grown for 15 hours at an average flow velocity *U*_gr_ = 2.10 mm*/*s. The two tests performed on each sample are conducted with initial flow velocities *U*_0_ = *U*_gr_ = 2.10 mm*/*s (stars) and *U*_0_ = *U*_gr_*/*5 = 0.42 mm*/*s (diamonds). The corresponding prestress values are *σ*_0_ = ⟨*σ*(*x*; *U*_gr_)⟩_12_ and *σ*_0_ = ⟨*σ*(*x*; *U*_gr_*/*5)⟩_12_. Open markers correspond to data measured first at *U*_0_ = *U*_gr_ and then at *U*_0_ = *U*_gr_*/*5, while closed markers are measured at *U*_0_ = *U*_gr_*/*5 and then at *U*_0_ = *U*_gr_. The dashed grey lines in **a**-**c** and **d**-**f** are the power law fits of the data reported in Fig. 2a and b, respectively. We verified that the observed strain-hardening behaviour is not significantly altered if fluid-structure interaction is accounted for in the calculation of the streamer rheology (Supplementary Fig. 9).

Given the dependence of viscoelastic properties on prestress, biofilm streamers should be compared based on tests conducted at a fixed prestress rather than at a fixed flow velocity. This is demonstrated by the PA14 wild-type and Δpel streamers, which show no significant differences in the differential Young’s modulus and effective viscosity when measured as a function of prestress but show apparent differences when results are aggregated by flow conditions (Supplementary Figure 5b, c) [2, 3].

### Stress-hardening behaviour in streamers across several bacterial species

Experiments conducted with several widely spread bacterial species, which are common infection agents and model organisms for biofilm research, demonstrate that stress-hardening is a universal feature of the biofilm streamer EPS matrix. In these experiments, we grow streamers using suspensions of *P. aeruginosa* PAO1, *B. cenocepacia* H111, *S. epidermidis* 1457, and *E. coli* RP437, flowing at *U*_gr_ = 2.10 mm*/*s for 15 h. These species differ in shape, motility and EPS components, with *S. epidermidis* being gram-positive and the others gram-negative bacteria [47–51]. Despite these differences, all species form biofilm streamers with an eDNA backbone, morphologically similar to those produced by PA14, as demonstrated by the strong fluorescence signal emitted by the PI-stained samples (Fig. 4a-d).

**Fig. 4.**
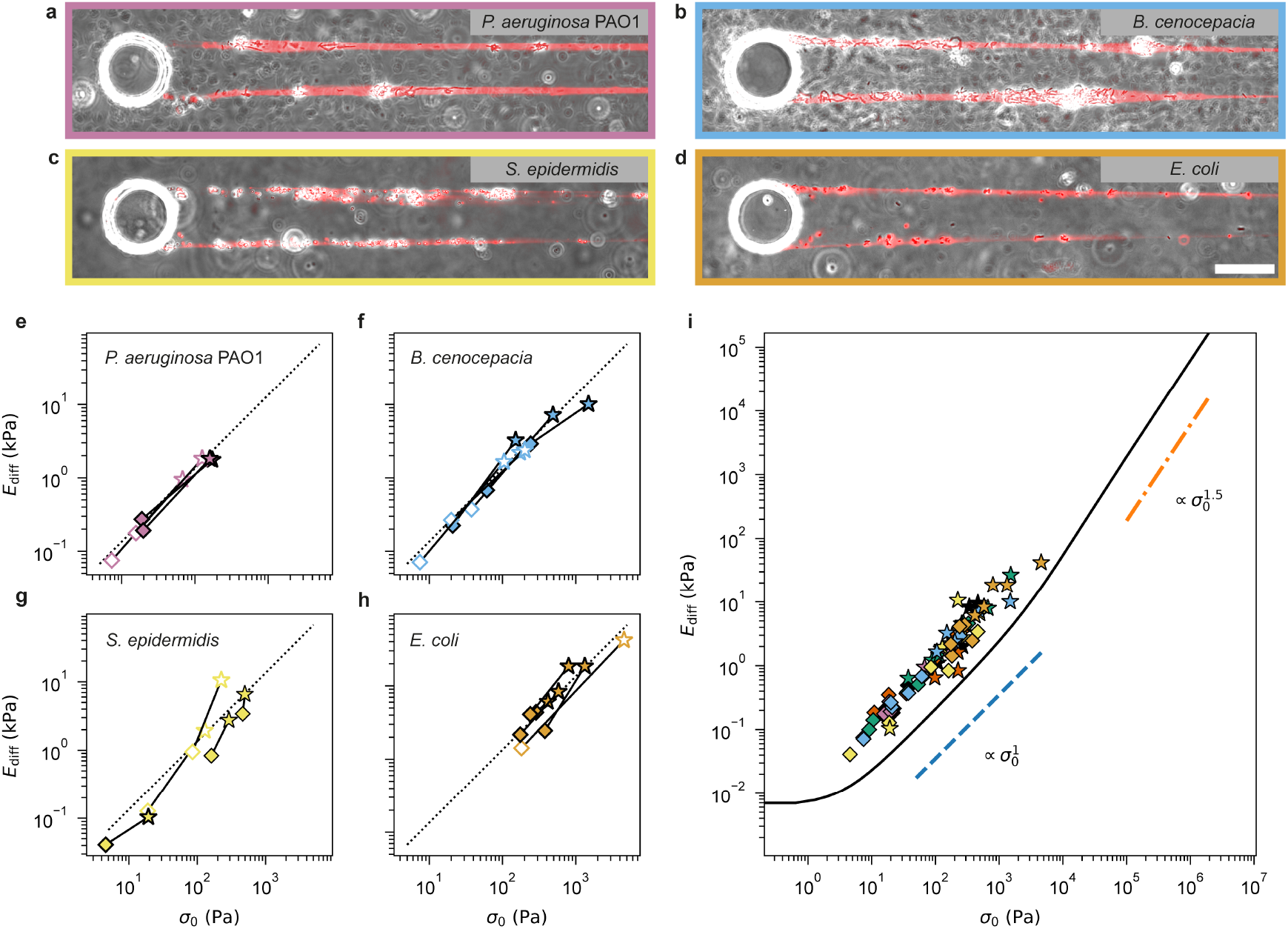
The stress-hardening behaviour of the EPS matrix is consistent across biofilm streamers from several bacterial species. **a**-**d**, Representative overlaid phase contrast and fluorescence images of portions of biofilm streamers grown for 15 hours at an average ambient flow velocity of *U*_gr_ = 2.10 mm*/*s by *P. aeruginosa* PAO1 (**a**), *B. cenocepacia H111* (**b**), *S. epidermidis* 1457 (**c**), and *E. coli* RP437 (**d**). The scale bar is 50 µm. **e**-**h**, Differential Young’s moduli *E*_diff_, obtained from sequential mechanical tests, plotted as a function of the axial stress *σ*_0_, demonstrating the stress-hardening behaviour of biofilm streamers formed by *P. aeruginosa* PAO1 (**e**), *B. cenocepacia* H111 (**f**), *S. epidermidis* 1457 (**g**), and *E. coli* RP437 (**h**), grown at average flow velocity *U*_gr_ = 2.10 mm*/*s. Solid lines connect results from the same portion of streamer tested under two axial stress conditions *σ*_0_. Tests are conducted on 15-hour-old streamers with initial flow velocities *U*_0_ = *U*_gr_ = 2.10 mm*/*s (stars) and *U*_0_ = *U*_gr_*/*5 = 0.42 mm*/*s (diamonds). The corresponding prestress values are *σ*_0_ = ⟨*σ*(*x*; *U*_gr_)⟩_12_ and *σ*_0_ = ⟨*σ*(*x*; *U*_gr_*/*5)⟩_12_. Open markers indicate data measured first at the higher and then lower values of *σ*_0_, while closed markers are measured at the lower and then at the higher values of *σ*_0_. The dotted lines in panels **e**-**h** indicate a power law behaviour with exponent equal to 1. **i**, The black line shows the differential Young’s modulus *E*_diff_ as a function of prestress *σ*_0_ for a 3D network of worm-like chains under uniaxial extension, modelled using the eight-chain model [52]. The curve was calculated as explained in the Supplementary Information, assuming *T* = 296 K, *A* = *l* = 50 nm, *N* = 1000, *n* = 10^21^ *m*^*−*3^. The corresponding contour length between neighbouring cross-links is *L*_c_ = *Nl* = 50 µm. The scaling in the crossover regime at intermediate prestress, where 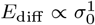, and in the high prestress regime, where 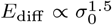, are indicated by dashed blue and dash-dotted orange lines, respectively. The markers show the data reported in Fig. 3a-c and panels **e**-**h** of this figure (same symbols and colour code of the original panels).

Sequential mechanical tests performed under different prestress conditions reveal the same stress-hardening behaviour observed in PA14 streamers (Fig. 4e-h). Specifically, the differential Young’s modulus of the streamers from all species scales linearly across a wide range of prestress, spanning nearly three orders of magnitude (Fig. 4i). Moreover, the prefactor also remains consistent across the testing conditions (Fig. 2 and Fig. 3) and different bacterial species (Fig. 4). This finding supports our conclusion that stress-hardening is independent of the specific composition of the biofilm matrix. In particular, the bacterial species tested produce different polysaccharides, yet all exhibited similar mechanical responses. The consistent stress-hardening behaviour and linear scaling observed across different species, combined with the presence of eDNA as a common and abundant component in their EPS matrices, suggests a link between this stress-hardening and the mechanical properties of the eDNA backbone.

### Entropic elasticity as the origin of stress-hardening behaviour in streamers

The stress-hardening behaviour of the differential Young’s modulus of the streamers can be attributed to the entropic elasticity of their eDNA backbone. We demonstrate this by modelling the expected elastic response of a DNA network under uniaxial extension, using a framework based on the worm-like chain model [52], and the persistence length of DNA [5] (Supplementary Information). This model accounts for the entropic elasticity of DNA molecules and was previously applied to describe the strain-hardening behaviour in DNA gels under shear [5, 53]. In the uniaxial extension configuration tested in our experiments, the model predicts three regimes: a linear elastic regime at low prestress, where *E*_diff_ is constant, a crossover regime at intermediate prestress where *E*_diff_ ∝*σ*_0_, and a regime at large prestress where 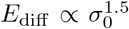 (Fig. 4i) (Supplementary Information). The observed agreement between the experimental and theoretical scaling exponents, which is commonly used as a robust indicator of the physical mechanism underlying stress-hardening in soft biological networks [43, 46, 54], supports the conclusion that the stress-hardening behaviour of streamers originates from the entropic elasticity of their eDNA backbone. This predicted crossover regime coincides with the prestress range probed in our experiments when assuming chain density *n ≈* 10^21^ *m*^*−*3^ and a contour length of DNA chains between neighbouring cross-links *L*_*c*_ = 50 µm (Fig. 4i). Parameter sweeps around the above values reveal that a broader set of (*n, N*) values yields a linear-scaling regime that fully spans – and exceeds – the prestress range explored experimentally (Supplementary Fig. 10, Fig. 4i). This suggests that the observed scaling does not require fine-tuning or specific assumptions about the microscopic configuration of the network, which remains to be characterised in detail. Instead, the scaling appears intrinsic to the underlying physics of the system and is likely to emerge under a wide range of conditions.

Although standard approaches for identifying the physical origin of stress-hardening typically do not require quantitative agreement between experimental and theoretical absolute values [43, 46, 54], the approximately fivefold discrepancy observed here prompts further investigation. In particular, we explore whether other EPS components, common across bacterial species, may contribute both to the stress-hardening behaviour and to the observed magnitude of the differential elastic modulus. To this end, we grow streamers and perform enzymatic treatments targeting extracellular RNA (eRNA) and extracellular proteins, with particular attention to the DNABII family of DNA-binding proteins, which are important EPS components conserved across species.

Degradation of eRNA with a mixture of RNases alters both the structure and mechanical response of the streamers (Fig. 5). Morphologically, RNase treatment leads to a reduced streamer radius and increased matrix density (Fig. 5a-f). This densification is so pronounced that the matrix becomes visible even in phase contrast images (Fig. 5f). Rheological tests show that eRNA degradation suppresses stress-hardening, as elasticity becomes independent of prestress in the prestress range tested (Fig. 5g). This behaviour is consistent with our model, which predicts that increasing polymer density shifts the onset of nonlinear elasticity to higher prestresses and raises the linear-elasticity plateau (Fig. 5g). These findings suggest that, although not essential for streamer integrity, eRNA modulates the structure of the matrix network, likely by influencing mesh size and crosslinking, and thereby alters its overall rheology.

**Fig. 5.**
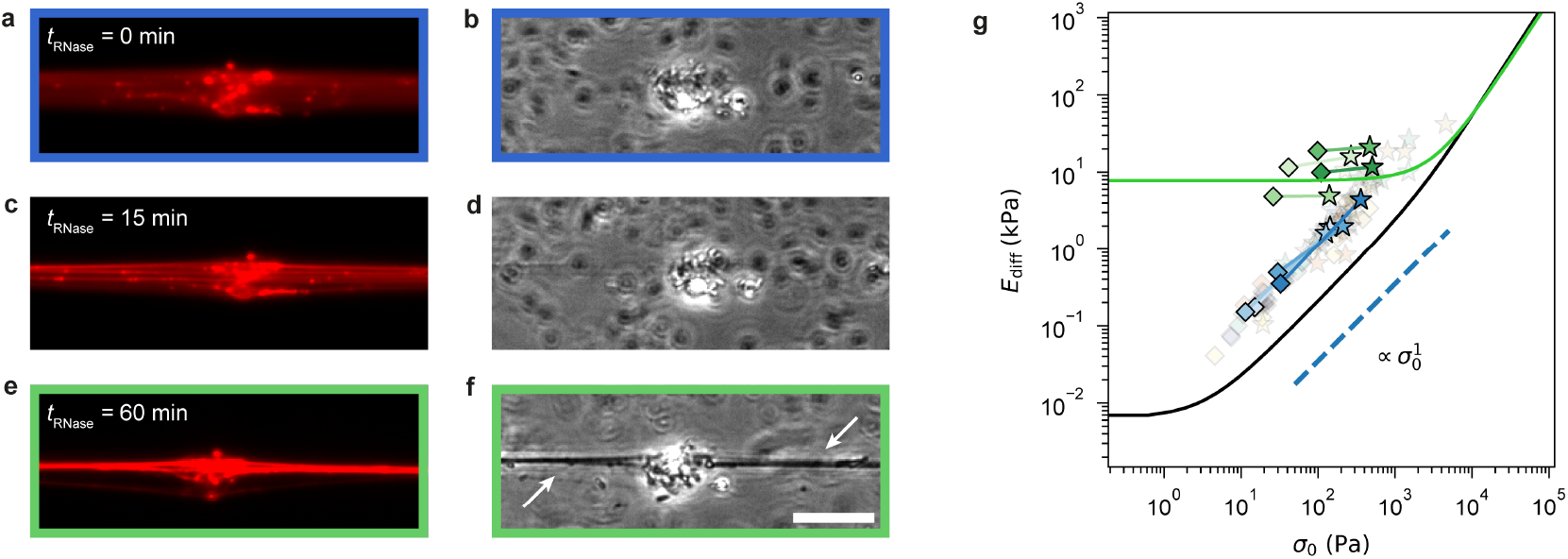
RNase treatment increases matrix density and suppresses stress-hardening within the accessible prestress range. **a**-**f**, Representative (**a, c, e**) fluorescence and (**b, d, f**) phase contrast images of a *P. aeruginosa* PA14 biofilm streamer (**a, b**) before (*t*_RNase_ = 0 min), (**c, d**) during (*t*_RNase_ = 15 min) and (**e, f**) after (*t*_RNase_ = 60 min) RNase treatment. RNase causes a reduction in streamer radius and an increase in matrix density. The densified matrix becomes visible even in phase contrast images (f, white arrows). Experiments were conducted on streamers grown for 15 hours at an average ambient flow velocity of *U*_gr_ = 2.10 mm*/*s. The scale bar is 20 µm. **g**, Differential Young’s moduli *E*_diff_, obtained from sequential mechanical tests, plotted as a function of the axial stress *σ*_0_, before (blue markers) and after (green markers) RNase treatment. Solid lines connect results from the same portion of the streamer tested under two prestress conditions *σ*_0_. Tests are conducted on 15-hour-old streamers with initial flow velocities *U*_0_ = *U*_gr_ = 2.10 mm*/*s (stars) and *U*_0_ = *U*_gr_*/*5 = 0.42 mm*/*s (diamonds), corresponding to *σ*_0_ = ⟨*σ*(*x*; *U*_gr_)⟩_12_ and *σ*_0_ = ⟨*σ*(*x*; *U*_gr_*/*5) ⟩_12_, respectively. Before RNase treatment, streamers show stress-hardening with scaling 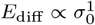 (dashed blue line), in agreement with previous results and with the model reported in Fig. 4i (black line). Semitransparent markers show the data of Fig. 4i, with no RNase treatment. After RNase treatment, elasticity becomes independent of prestress, indicating entry into the linear-elastic plateau. Our theoretical model predicts that increasing network density shifts the onset of nonlinear elasticity to higher prestress values and raises the value of the linear-elastic plateau. Accordingly, the densification observed after RNase treatment likely shifts the prestress range accessible in our setup into this plateau.

In contrast, proteinase K treatment has no observable effect on streamer morphology and stress-hardening (Supplementary Fig. 12), suggesting that proteins, including the ones belonging to the DNABII-family, do not play significant structural or mechanical roles in this system.

Taken together, our findings align with recent studies on eNA networks extracted from surface-attached *P. aeruginosa* biofilms, which showed that neither polysaccharides nor proteins are required for the stability of eDNA network in the biofilm matrix [41]. This agreement supports the conclusion that eNA are the primary structural and mechanical component of streamers, and that their interactions modulate the observed entropic stress-hardening behaviour.

## Conclusion

Although the colonisation success of biofilms in dynamic and mechanically challenging habitats has long suggested the presence of adaptive mechanical responses enhancing their resilience, direct evidence has been lacking. Here, we demonstrate a strong adaptation of biofilm streamer viscoelasticity to varying mechanical stress conditions, driven by the stress-hardening behaviour of the biofilm matrix. Stress-hardening not only explains the ability of streamers to withstand rapid mechanical perturbations but also accounts for the difference in viscoelastic properties measured in samples grown under different flow velocities. Remarkably, this behaviour is consistent across multiple bacterial species, reflected in a linear dependence of the differential Young’s modulus and extensional viscosity on the axial prestress exerted by fluid flow. This stress-hardening response is observed across a wide range of prestress values, spanning three orders of magnitude, suggesting its role as a highly effective adaptation mechanism in a broad range of environments.

The experimentally observed linear scaling between differential Young’s modulus and prestress is consistent with the predicted elastic response of a worm-like chain network under uniaxial extension, based on the force-extension relation characteristic of DNA molecules [31, 52]. This indicates that the stress-hardening behaviour of streamers is entropic in nature and governed by the elasticity of the eDNA matrix. Notably, while the stress-hardening response is independent of the presence and type of polysaccharides and proteins within the matrix, our experiments show that eRNA modulates both the morphology and mechanical behaviour of biofilm streamers. These findings support the conclusion that the mechanical response is primarily associated with the eNA backbone, which is conserved across all tested species, as visualized using PI staining.

eDNA molecules are fundamental components of streamers, essential for their integrity [3] and ultimately responsible for their stress-hardening behavior. Together with the results from enzymatic RNase treatment, this suggests that the backbone of streamers is formed by eDNA molecules. Their cross-links could consist of Holliday junctions, branched structures generated by the partial exchange of strands between two double-stranded DNA molecules. Such junctions have been observed in the matrix of in vivo biofilms, with inter-junction distances the order of ~ 1–10 µm [33], which we suggest may be compatible with the ~ 50 µm contour length between neighbouring cross-links inferred from our modelling results (Fig. 4i), supporting their potential role in stabilizing the eDNA network in streamers. Considering the mechanical behaviour of DNA itself, its entropic elasticity and strain-stiffening are well documented both at the single-molecule level [5, 31, 55–57] and in synthetic gels [53]. However, our study provides the first direct evidence that this intrinsic mechanical property of DNA contributes functionally to the mechanics of a naturally assembled biological network as the biofilm matrix.

Moreover, enzymatic RNase treatment of streamers suggests that eRNA plays a secondary role: while not essential for their integrity, it modulates network elasticity, likely by altering molecular and cross-link density in the matrix. Specifically, the removal of eRNA leads to a visible densification of the biofilm matrix and a marked shift in mechanical behaviour: elasticity becomes independent of prestress, and the stress-hardening response is suppressed in the probed prestress range. This change is consistent with the worm-like chain network model, in which increased matrix density shifts the onset of nonlinearity and raises the linear-elastic plateau. These modelling results support the hypothesis that the removal of eRNA frees up space or bonding sites for tighter crosslinking of eDNA, thereby increasing matrix stiffness. However, a full experimental characterisation of the eNA network microstructure of streamers and the development of a more detailed theoretical model will be required to mechanistically account for the role of eRNA, and not just eDNA, in the matrix behaviour.

Our experiments show a remarkable functional role of NA in their extracellular state. The purely mechanical adaptation mechanism we uncovered allows streamers to maintain cohesion under varying stress conditions, with potential implications for biofilm structural integrity, ecological resilience, and colonisation dynamics. Stress-hardening preserves matrix integrity by increasing stiffness under high stress, while allowing greater deformability in low-stress conditions, thereby enhancing adaptability in fluctuating environments. In our experiments, the increase in differential Young’s modulus with prestress consistently limits the elastic strain increment to below 10% across all conditions (Supplementary Fig. 5a). By limiting deformation and maintaining bacterial community cohesion under mechanical load, stress-hardening may offer an ecological benefit in dynamic ecosystems, helping to keep community members within interaction distance [58]. Furthermore, we propose that the stress-hardening behaviour observed in biofilm streamers may have far-reaching implications for biofilm development and survival, paralleling its role in biological tissues, where it helps preserve structural integrity and regulate key processes such as morphogenesis and wound healing [59, 60]. Given that eNA are widespread components of biofilms with different architectures [1, 6, 41], this mechanical response may extend well beyond the specific case of streamers. While the biological and ecological consequences of stress-hardening in biofilms remain to be fully elucidated, our study provides direct *in situ* evidence that eNA contribute both structurally and functionally to the mechanical adaptability of the biofilm matrix, laying the groundwork for future investigations into the physical strategies underpinning microbial community organisation and persistence.

## Methods

### Bacterial culture conditions

The following bacterial species are used in this study: *Pseudomonas aeruginosa* PA14 wild-type, the Pel deletion mutant PA14 ΔpelE (referred to as Δpel in the manuscript), the Pel-overproducer PA14 ΔwspF, *Pseudomonas aeruginosa* PAO1 wild-type, *Burkholderia cenocepacia* H111, *Staphylococcus epidermidis* 1457 and *Escherichia coli* RP437. The *B. cenocepacia* suspensions are prepared according to [30] by inoculating cells from a frozen stock in 3 ml of ABC medium and incubating overnight at *T* = 37 °C while shaking at 200 rpm. 30 µl of the overnight suspension are then inoculated in 3 ml of ABC medium and incubated for 6 hours at *T* = 37 °C while shaking at 200 rpm, until OD_600_ = 0.3 is reached. The overnight suspensions are then diluted in fresh Tryptone Broth (10 g/l tryptone, 5 g/l NaCl) to a final optical density OD_600_ = 0.006 and loaded into glass syringes. For all other bacterial species, suspensions are prepared by inoculating Luria

broth 1.5% agar plates from frozen stocks and incubating them overnight at *T* = 37 °C. Cells from a single colony are inoculated in 3 ml of liquid medium and incubated at *T* = 37 °C for 3 hours while shaking at 200 rpm, until OD_600_ = 0.3 is reached. M9 minimal medium (0.4% glucose) is used for *E. coli*, and Tryptone Broth (10 g/l tryptone, 5 g/l NaCl) is used for all other strains. The suspensions are then diluted in the respective fresh liquid media to a final optical density of OD_600_ = 0.006 and loaded into glass syringes.

### Formation of biofilm streamers under different ambient flow conditions

Biofilm streamers are cultured and tested in the polydimethylsiloxane (PDMS) microfluidic platform described in [2]. Briefly, the platform consists of four straight channels with isolated micropillars that act as nucleation sites for pairs of biofilm streamers when a bacterial suspension is flowed at a constant average flow velocity *U*_gr_. The channels have width *W* = 1 mm, height *H* = 40 µm and length *L* = 4 cm. The cylindrical micropillars span the whole height of the channel, and their radius is *R*_pillar_ = 25 µm. In this study, the number and locations of the micropillars within each microchannel are slightly modified from those of the original platform to enhance the efficiency of streamer formation. In the new configuration, five pillars are positioned at varying *y*-coordinates across the width of each channel (Supplementary Fig. 7c), rather than being aligned at the channel half-width as in the original configuration. Such a staggered arrangement ensures that the pillars intercept different flow lines without screening each other from the biomass in flow. Four laminar flow conditions are tested in parallel across different channels of the same microfluidic platform. The same bacterial batch is used in all channels to minimise biological variability. All streamers in this study are grown at room temperature (*T* = 23 *±* 1 °C). The average flow velocities *U*_gr_ span an order of magnitude, corresponding to Reynolds numbers Re ranging from 0.02 to 0.20 (Supplementary Table 1). The Reynolds number is calculated as Re = *ρU*_gr_*D/µ*, where *ρ* is the density of water, *D* is the pillar diameter, and *µ* is the dynamic viscosity of water. Flow velocities are controlled using a syringe pump (neMESYS 290N, CETONI, Germany). To maintain a constant flux of cells across the four different flow conditions, two syringes are used for each microfluidic channel (Supplementary Fig. 7a): one to inject bacterial suspensions at a constant flow rate *Q*_cells_ = 0.06 ml*/h* and the other to inject growth medium at a flow rate *Q*_medium_. The value of *Q*_medium_ is set so as to reach the desired total flow rate *Q*_gr_ = *Q*_cells_ + *Q*_medium_ in each channel. In the channel with the lowest flow rate, only the first syringe is used, such that *Q*_gr_ = *Q*_cells_. The values of *Q*_gr_ used in this study correspond to the average flow velocities *U*_gr_ = *Q/*(*HW*) reported in Supplementary Table 1. The streams from the two syringes are joined by a Y connector (P-514, IDEX) located before the channel inlet. After the Y connector, a 2.5-mm-wide and 2.2-cm-long straight connector (from the LVF-KMM-06 kit, Darwin Microfluidics) is added to ensure complete sample homogenisation before entering the channel. In the *Wisteria floribunda* lectin staining experiments, a third syringe is added to introduce the lectin solution once the streamers are formed (Supplementary Fig. 7b). This is achieved by incorporating an additional Y connector (P-514, IDEX) before the channel inlet. A shut-off valve (P-732, IDEX) is mounted on the lectin flow line to allow the delayed mounting of the lectin syringe.

## Characterisation of streamer composition and morphology

Streamer composition and morphology are characterised as described in [2]. Briefly, the biofilm matrix components are fluorescently stained and then imaged using epifluorescence microscopy. eDNA in the matrix is stained using propidium iodide (PI), which is added to both the bacterial suspension and the growth medium syringes to a final concentration of 2 µg*/*ml.

The red fluorescence signal of PI is acquired at half-height of the microchannel, and then binarised and processed to determine the morphology of the streamers according to [2]. Specifically, image processing provides the radius *R*(*x*) and centre *C*(*x, y*_c_(*x*)) of the streamers’ cross sections as functions of the streamwise coordinate *x*. This data is then used to measure the streamers’ lengths *L* and average radii ⟨*R*⟩ over ROI_1_ (*x* ∈ [25 µm, 125 µm]) and ROI_2_ (*x* ∈ [400 µm, 1065 µm]). Data reported in Fig. 1e, f are the values of *L* and ⟨*R*⟩ averaged over about 10 experimental replicates for each strain and flow condition. The morphological characterisation from the PI signal, acquired right before each mechanical test, is also used to build 3D models of the streamers for the numerical calculation of the corresponding axial prestresses *σ*_0_, following the protocol presented in [2]. Additionally, the average eDNA intensity reported in Supplementary Fig. 4 is calculated for each sample by averaging the PI signal between the initial positions *x*_1_, *x*_2_ of the cell aggregates tracked during the rheological tests as

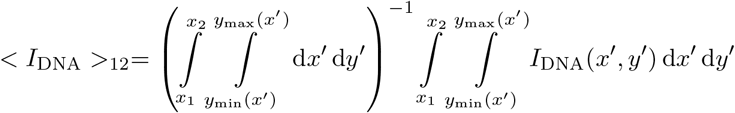

where *y*_min_(*x*) = *C*(*x, y*_*c*_(*x*)) −*R*(*x*)) and *y*_max_(*x*) = *C*(*x, y*_*c*_(*x*)) + *R*(*x*)). The integrals were performed as discrete sums over the image pixels. Pel in the matrix of PA14 wild-type and ΔwspF was stained with GFP fluorescently labelled *Wisteria floribunda* lectin (WFL, FL-1351-2, Vector Laboratories). WFL is diluted in phosphate-buffered saline solution at 10 µg*/*ml concentration and loaded into a syringe. Endpoint measurements are performed on 15-hour-old biofilm streamers by flowing WFL in the microchannels for 30 min before image acquisition in the GFP fluorescence channel. The WFL syringes are mounted after 15 hours from the start of the experiment to avoid overnight degradation of WFL. Tests for quantifying Pel abundance with WFL are performed on streamers grown without PI to avoid interactions between the two stains. After imaging WFL, the WFL syringes are replaced with PI syringes to stain and quantify eDNA abundance. Phase contrast microscopy is used to image bacterial cells. The experiments are performed at ambient temperature (*T* = 23 *±* 1 °C).

### Characterisation of streamer growth

Streamer growth under different flow conditions *U*_gr_ is characterised using 24-hour-long time-lapse acquisitions. Phase contrast and mCherry fluorescence images are acquired every 15 minutes. The length *L* as a function of time, is measured by tracking the streamer tip on the mCherry fluorescence channel. Analysis of these time series showed that the initial growth stages often involved significant detachment events. These events are defined as reductions in length *L* below a minimum threshold of *L*_0_ = 250 µm. At later stages, streamers typically reach a more stable growth regime. The onset of this second regime is identified for each streamer by the time *t*_0_ at which *L*(*t*) *> L*_0_ for all subsequent time points *t > t*_0_. The average growth speed as a function of time for the second regime (Supplementary Fig. 1d) is calculated as the derivative of the power law fit reported in Supplementary Fig. 1c. The length threshold *L*_0_ is set to a value corresponding to a distance 10*R*_pillar_ from the pillar. At this distance, the flow velocity magnitude matches its unperturbed value far from the pillar, with a relative difference of less than 0.1% (Supplementary Fig. 1a). This choice allows us to neglect phenomena occurring near the pillar, where biofilm-substrate adhesion and hydrodynamic interaction with the complex flow field play an important role. The average waiting time *t*^*\**^ for the first irreversible attachment event of dead cells or eDNA aggregates to the pillar is calculated from the PI signal on the pillar surface (Supplementary Fig. 1b). The waiting time in each experimental replicate is measured as the time at which the first irreversible attachment event of PI-stained biomass on the pillar is observed, with a minimum size threshold set by the approximate size of a single bacterial cell. Data reported in Supplementary Fig. 1b, c for *t*_0_, *t*^*\**^ and *L − L*(*t*_0_) are averaged over about 15 streamers for each flow condition.

### Rheological tests

The rheological characterisation of biofilm streamers grown under different ambient flow conditions is performed *in situ* with mechanical tests following the technique described in [2]. For each flow condition *U*_gr_, the mechanical tests are conducted imposing a three-stage flow profile: an initial 2.5-minute stage at flow velocity *U*_0_ = *U*_gr_, a 5-minute creep stage at *U*_cr_ = 2*U*_0_ = 2*U*_gr_, and a final 2.5-minute recovery stage at *U*_rec_ = *U*_0_ = *U*_gr_ (Supplementary Fig. 2a). The initial axial stress *σ* (*x*; *U*_gr_) in each test is numerically calculated with a computational fluid dynamics (CFD) simulation (COMSOL Multiphysics [61]) solving the Navier-Stokes equations of flow at average velocity *U*_gr_ around a 3D model of the streamer. The model is built based on the morphology determined by the PI signal acquired before the beginning of the test at average flow velocity *U*_gr_, as described in [2]. During the test, we acquire phase contrast images of the streamer in ROI_2_ at 1 fps. The deformation of a portion of streamer delimited by two cell aggregates located at *x*_1_ and *x*_2_ is measured by tracking their relative distance *l*_12_ (*t*) = *x*_2_ (*t*) *− x*_1_ (*t*) over time, providing the strain increment Δ*ε* (*t*) = (*l*_12_ (*t*) *− l*_12_ (0))*/l*_12_ (0) (Supplementary Fig. 2c). The typical strain increment is well described by a Maxwell model, with an instantaneous elastic contribution Δ*ε*_el_ and a viscous deformation Δ*ε*_visc_ (*t*) = ε̇ *t* linearly dependent on time, where ε̇ is the strain rate. The stress values averaged over the examined portion of streamer (*x* ∈ [*x*_1_, *x*_2_]) during the three stages of the test are: the prestress *σ*_0_ ≡ ⟨*σ* (*x*; *U*_0_)⟩_12_ during the initial stage; the creep stress *σ*_cr_ ≡ ⟨*σ* (*x*; *U*_cr_)⟩_12_, during the creep stage; and the recovery stress *σ*_rec_ = *σ*_0_ (Supplementary Fig. 2b). The notation ⟨*·*⟩_12_ denotes the average between *x*_1_ and *x*_2_. The applied extensional stress increment is Δ*σ* = *σ*_cr_ *− σ*_0_. Assuming negligible fluid-structure interaction, *σ*_rec_ = *σ*_0_ and the stress increment is Δ*σ* = *σ*_0_. Thus, the numerical estimate of *σ*_0_ and the measurements of Δ*ε*_el_ and ε̇ allow the calculation of the differential Young’s modulus *E*_diff_ = Δ*σ/*Δ*ε*_el_ and effective viscosity *η* = Δσ/ε̇ of the examined portion of streamer as a function of the applied extensional prestress *σ*_0_.

To obtain meaningful estimates of *E*_diff_ and *η*, Δ*σ* has to be selected such that the sample’s response is locally linear in a neighbourhood of *σ*_0_. We verified that the choice Δ*σ* = *σ*_0_ satisfies this condition (Supplementary Fig. 8).

To test the stress-hardening behaviour of the matrix, streamers are grown at *U*_gr_ = 2.10 mm*/*s for 15 hours. Each sample undergoes two sequential mechanical tests, each at a different initial flow velocity *U*_0_ and corresponding creep velocity *U*_cr_ = 2*U*_0_. The first test is performed at *U*_0_ = *U*_gr_ = 2.10 mm*/*s, corresponding to a prestress *σ*_0_ = ⟨*σ*(*x*; *U*_gr_)⟩_12_. After the first test, the flow velocity is decreased to *U*_gr_*/*5 = 0.42 mm*/*s, and the system is allowed to stabilise for 10 minutes. The second test is then performed at *U*_0_ = *U*_gr_*/*5 = 0.42 mm*/*s, corresponding to a presetress *σ*_0_ = ⟨*σ*(*x*; *U*_gr_*/*5)⟩_12_ *<* ⟨*σ*(*x*; *U*_gr_)⟩_12_ (Supplementary Fig. 2d). Before each test, the PI signal is acquired to characterise the morphology of the streamer at the corresponding *U*_0_. These morphologies are then used to numerically calculate the corresponding axial stresses with CFD simulations, according to [2]. During two sequential tests on the same streamer, the relative distance *l*_12_ (*t*) between the same pair of cell aggregates is tracked. This allows the quantification of *E*_diff_ and *η* of the same portion of streamer at the two different prestress values.

To estimate the effect of fluid-structure interaction (FSI) on the correlation between *E*_diff_ and *σ*_0_ (Supplementary Fig. 9), we applied the iterative scheme presented in [2] to the dataset reported in Fig. 3b. The scheme is arrested at the first iteration since higher-order iterations have a negligible effect on the FSI correction, as shown by data reported in [2].

### Proteinase K and RNase treatments

To test the role of proteins and RNA in streamer structure and viscoelasticity, PA14 wild-type streamers are grown at *U*_gr_ = 2.10 mm*/*s for 15 hours and then treated for 1 hour with proteinase K (Proteinase K, 1.24568, Sigma-Aldrich) or a mixture of RNases (RNase, 10109134001, Roche) at *T* = 23 *±* 1 °C and *T* = 37 °C, respectively. The temperature during treatment is controlled with a microscope stage-top incubator (H301-NIKON-NZ100/200/500-N chamber and H401-T-CONTROLLER temperature controller, Okolab). Proteinase K is dissolved in tryptone broth at a concentration of 1 mg*/*ml, and RNase is dissolved in 50 mM Tris-HCl buffer (pH 7.5) at a concentration of 1 mg*/*ml. Enzyme solutions are prepared and loaded into glass syringes after streamer growth to prevent enzyme degradation overnight. To assess the dependence of the differential Young’s modulus *E*_diff_ on prestress *σ*_0_ after treatment, each sample undergoes two sequential mechanical tests as described in the previous paragraph, at room temperature (*T* = 23 *±* 1 °C). Control experiments are performed at room temperature after treatment with RNase buffer alone at *T* = 37 °C with RNase buffer only (Supplementary Fig. 11).

## Data availability

The datasets generated and analysed during the current study are available from the corresponding author on reasonable request. Source Data are provided with this paper.

---

**Supplementary Fig. 1.**
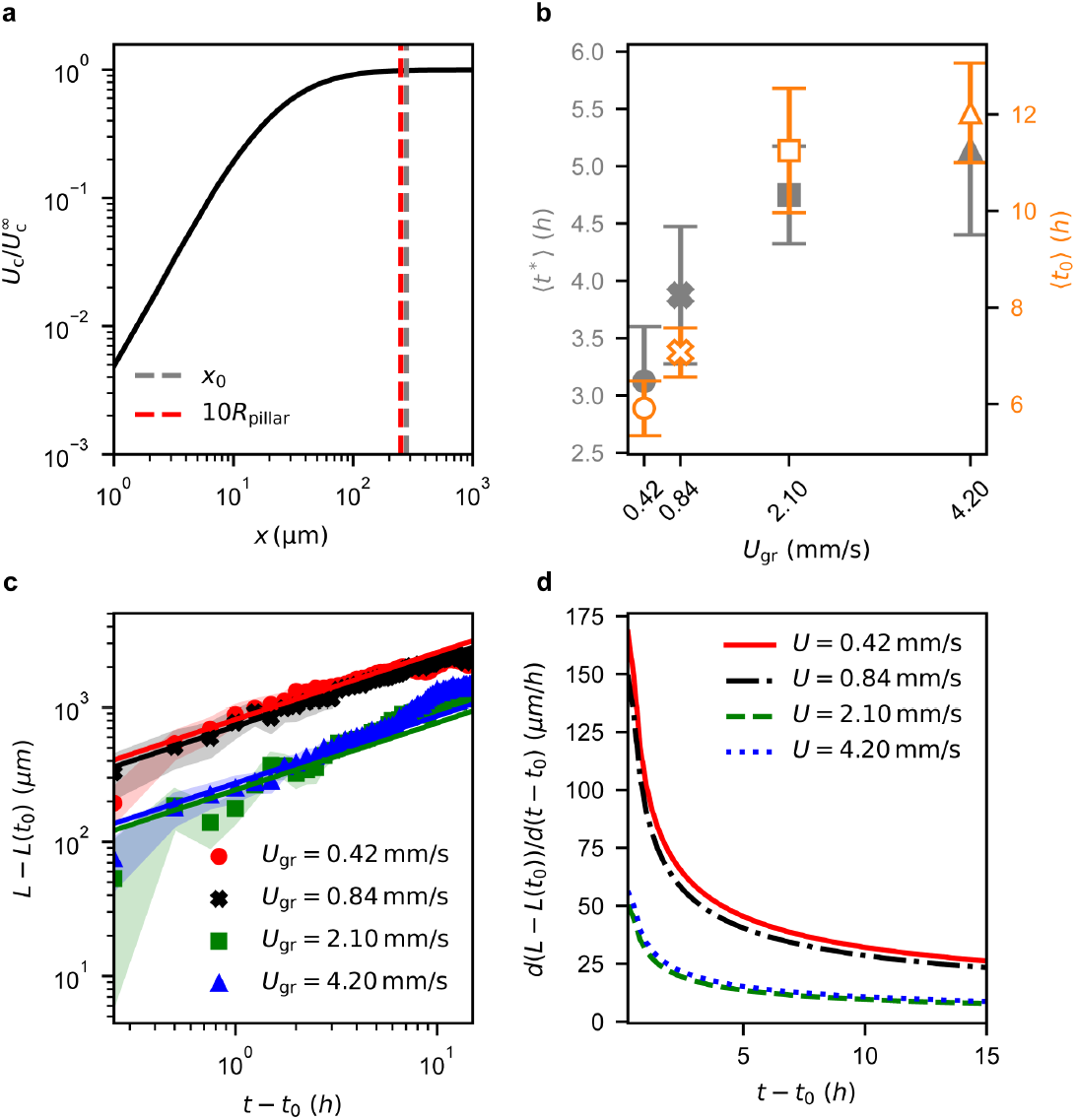
Effects of flow on the time scale of streamer inception and growth speed. **a**, Flow velocity magnitude *U*_c_ on the centerline of the microchannel as a function of the distance *x* from the pillar in the absence of streamers. The flow field was numerically calculated with a 3D CFD simulation of a pillar located at half-width of the channel. The dashed grey line indicates the distance *x*_0_ where the relative difference between *U*_c_ and the unperturbed value 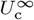 becomes less than 1%. The dashed red line shows that this position approximately corresponds to 10*R*_pillar_. **b**, Average waiting time *t*^*\**^ for the first irreversible attachment event of PI-stained cells or eDNA aggregates on the pillar (closed grey markers, left-hand *y*-axis) and time *t*_0_ needed to observe a stable filament growing without major detachment events defined as drops of *L* below *L*_0_ (open orange markers, right-hand *y*-axis). **c**, Averages of *L*(*t − t*_0_) *− L*(*t*_0_) as a function of time in the stable growth regime (*t >* −*t*_0_) for the four different flow conditions tested in this study. *L*(*t*) curves measured for single streamers were averaged from *t*_0_ onwards after subtracting the value *L*(*t*_0_). This method excluded the initial stages of streamer formation, where major detachment events occur. The solid lines show best fits with (*t − t*_0_)^0.5^. **d**, Average growth speeds as a function of time for the different flow conditions, obtained as the time derivative of the fitting curves in **c**. The average growth speeds at *t* = 15 h are lower than 50 µm*/*h in all flow conditions,. Therefore, streamer growth is negligible during the time scale of the rheological tests (~ 10 min).

**Supplementary Fig. 2.**
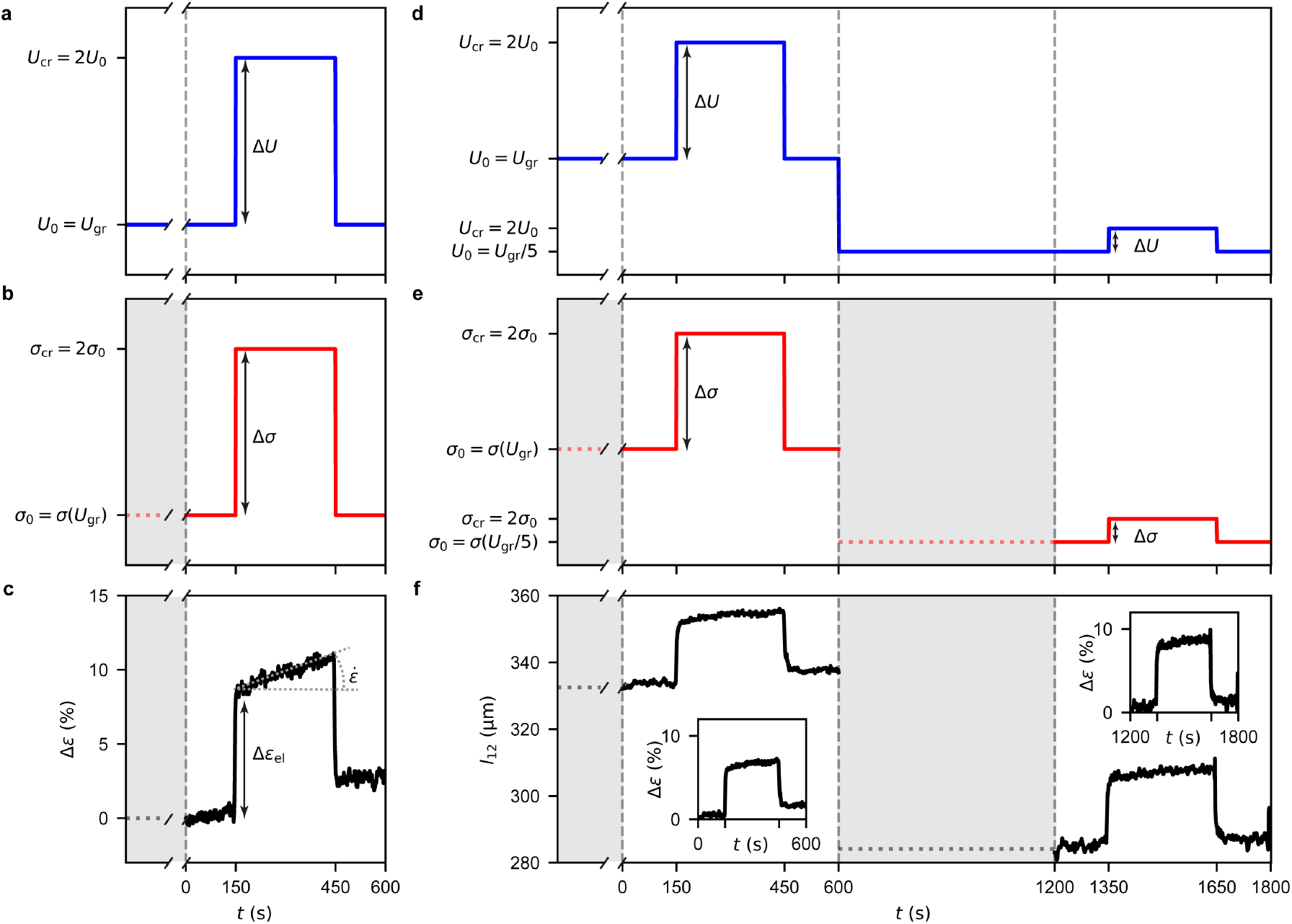
Mechanical test scheme. **a**-**c**, Average flow velocity *U*, axial stress *σ* and typical strain increment Δ*ε* of a streamer portion as a function of time *t* during the 10-minute mechanical tests for the characterisation of biofilm streamers grown under different ambient flow conditions *U*_gr_ (Fig. 2). **a**, In the 2.5-minute initial stage of the test, the flow velocity is *U*_0_ = *U*_gr_; in the 5-minute creep stage *U*_cr_ = 2*U*_0_ = 2*U*_gr_; in the 2.5-minute recovery stage, *U* is decreased to the initial value *U*_rec_ = *U*_0_ = *U*_gr_. The dimension line shows the velocity increment Δ*U* imposed during the creep stage. The broken axis shows that the initial flow velocity *U*_0_ equals the average flow velocity *U*_gr_, maintained constant during streamer formation. **b**, The prestress *σ*_0_ applied to the tested portion of the streamer during the initial stage corresponds to the axial stress exerted by flow at *U*_gr_: *σ*_0_ = ⟨*σ*(*x*; *U*_gr_)⟩_12_. The doubling of the flow velocity during the creep stage corresponds to doubling the axial stress *σ*_cr_ = 2*σ*_0_, if fluid-structure interaction is neglected. In the recovery stage of the test, the axial stress decreases to its initial value: *σ*_rec_ = *σ*_0_. The dimension line shows the stress increment Δ*σ* = *σ*_cr_ *− σ*_0_ imposed during the creep stage. The axial stress before the beginning of the test (shaded region) is not determined: *σ* (*x*; *U*_0_) was characterised with 3D CFD simulations based on the PI fluorescence signal acquired before the test, at *t* = 0. **c**, The strain increment Δ*ε* (*t*) of the examined portion of streamer is well described by a Maxwell model, with an instantaneous elastic contribution Δ*ε*_el_ at *t* = 150 s and an irreversible time-dependent viscous deformation with strain rate ε̇, which takes place during the creep stage of the test. When the axial stress increment is removed (*t* = 450 s), only the elastic deformation is recovered. The streamer portion is delimited by two cell aggregates, and its deformation is characterised by tracking their relative distance *l*_12_ (*t*) on phase contrast images acquired at 1 fps, yielding the strain increment as Δ*ε* (*t*) = (*l*_12_ (*t*) −*l*_12_ (0)) */l*_12_ (*t*). **d**-**f**, Average flow velocity *U*, axial stress *σ* and typical length *l*_12_ of a streamer portion as a function of time *t* during two sequential mechanical tests (Fig. 3, Fig. 4). **d**, After 15 hours of growth, a first 10-minute mechanical test like the one represented in panels **a**-**c** is performed on a streamer at *U*_0_ = *U*_gr_ (*t* ∈ [0, 600 s]). After the first test, the flow velocity is decreased to *U* = *U*_gr_*/*5. After waiting 10 minutes for stabilisation, a second 10-minute test is performed at *U*_0_ = *U*_gr_*/*5 (*t* ∈ [1200 s, 1800 s]). The corresponding average velocities during the creep and recovery stages during the second test are *U*_cr_ = 2*U*_0_ = 2*U*_gr_*/*5 and *U*_cr_ = *U*_0_ = *U*_gr_*/*5, respectively. **e**, Stress values applied to the same portion of streamer during the sequential tests. In the first and second tests, the applied prestresses are *σ*_0_ = ⟨*σ*(*x*; *U*_gr_)⟩_12_ and *σ*_0_ = ⟨*σ*(*U*_gr_*/*5)⟩_12_, respectively. The prestress values are numerically calculated with 3D CFD simulations based on the PI fluorescence signal acquired before each test, at *t* = 0 and *t* = 1200 s. Since the two morphologies differ, *σ*(*x*; *U*_gr_*/*5) *σ*(*x*; *U*_gr_)*/*5. **f**, Typical length *l*_12_ (*t*) of the portion of streamer delimited by the two cell aggregates tracked during the two tests, measured as explained in **c** during the first (*t* ∈ [0, 600 s]) and the second test (*t* ∈ [1000 s, 1800 s]). The insets show the corresponding strain increments Δ*ε*.

**Supplementary Fig. 3.**
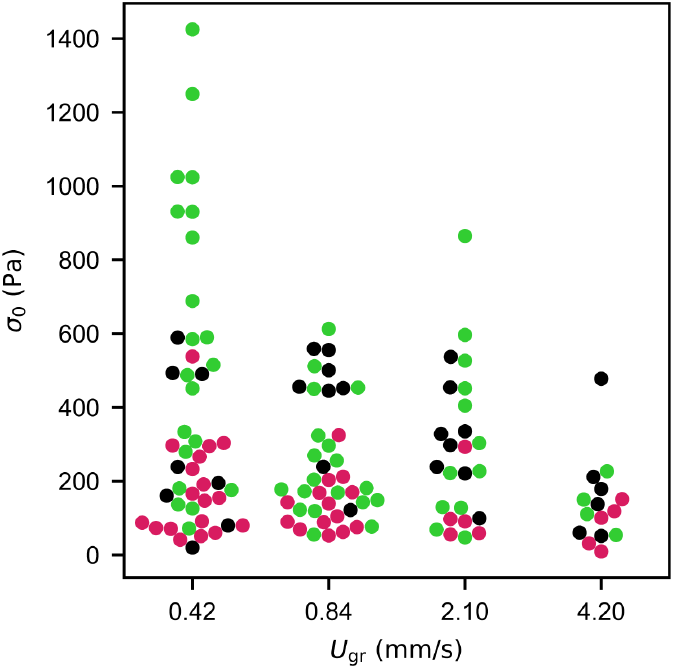
Distributions of prestress values *σ*_0_ corresponding to the flow velocities *U*_gr_ tested in this study. The swarm plot illustrates the axial prestress values *σ*_0_ exerted on the tested portion of each sample reported in Fig. 2. The axial stress corresponds to the prestress exerted by the background flow with average velocity *U*_gr_ on the examined portion of streamer during the 2.5-minute initial stage of the rheological test. The swarm plot reports data for PA14 Δpel (magenta), wild-type (black), and ΔwspF (green). Since *σ*_gr_ is determined both by *U*_gr_ and streamer morphology, while it is true that streamers grown at a higher ambient flow condition are subjected to higher shear stresses and pressure gradients, it cannot be assumed that they are also subjected to higher axial stresses. This is clearly demonstrated at *U*_gr_ = 4.20 mm*/*s, the maximum *U*_gr_ tested. Despite being associated with the highest average shear stress and pressure gradient values, it exhibits the lowest average *σ*_0_. This observation can be explained in light of the streamer morphology under this flow condition: lengths are shortest, and radii are widest (Fig. 1e-f), which decreases the overall axial stress on the streamer.

**Supplementary Fig. 4.**
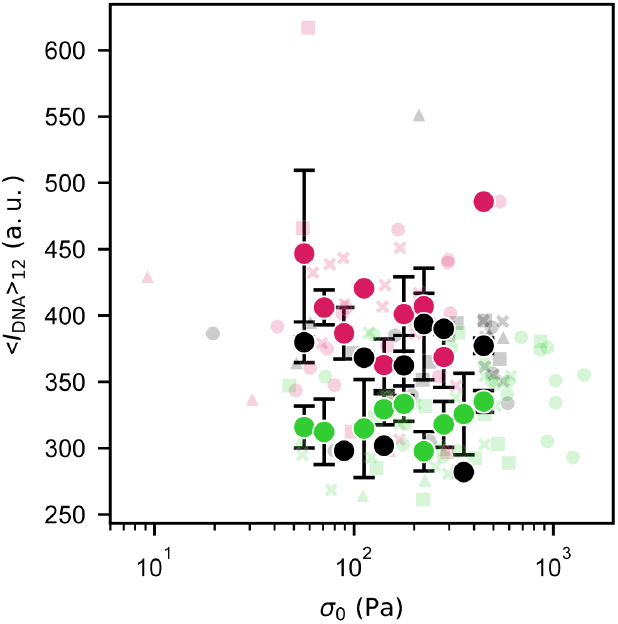
Average eDNA intensity *I*_DNA_ on the tested portion of streamer as a function of the applied prestress *σ*_0_. Semitransparent markers show the intensity of the eDNA fluorescence signal *I*_DNA_ averaged over the length of each tested portion of streamer reported in Fig. 2, as a function of the applied prestress *σ*_0_ = ⟨*σ* (*x*; *U*_gr_)⟩_12_. Each value of ⟨*I*_DNA_⟩_12_ was calculated between the positions *x*_1_ and *x*_2_ of the cell aggregates tracked during the stress test, as explained in the Methods. The different colours represent the different PA14 mutants: Δpel (magenta), wild-type (black) and ΔwspF (green). The different symbols represent data points obtained for samples grown at different average flow velocities *U*_gr_: 0.42 mm*/*s (circles), 0.84 mm*/*s (crosses), 2.10 mm*/*s (squares) and 4.20 mm*/*s (triangles). Fully opaque markers were calculated by binning and averaging the semi-transparent points for each bacterial strain within 10 intervals of *σ*_0_. These bins have equal width on a logarithmic scale and cover the stress range where data was collected for all the mutants. The error bars show the standard error of the mean.

**Supplementary Fig. 5.**
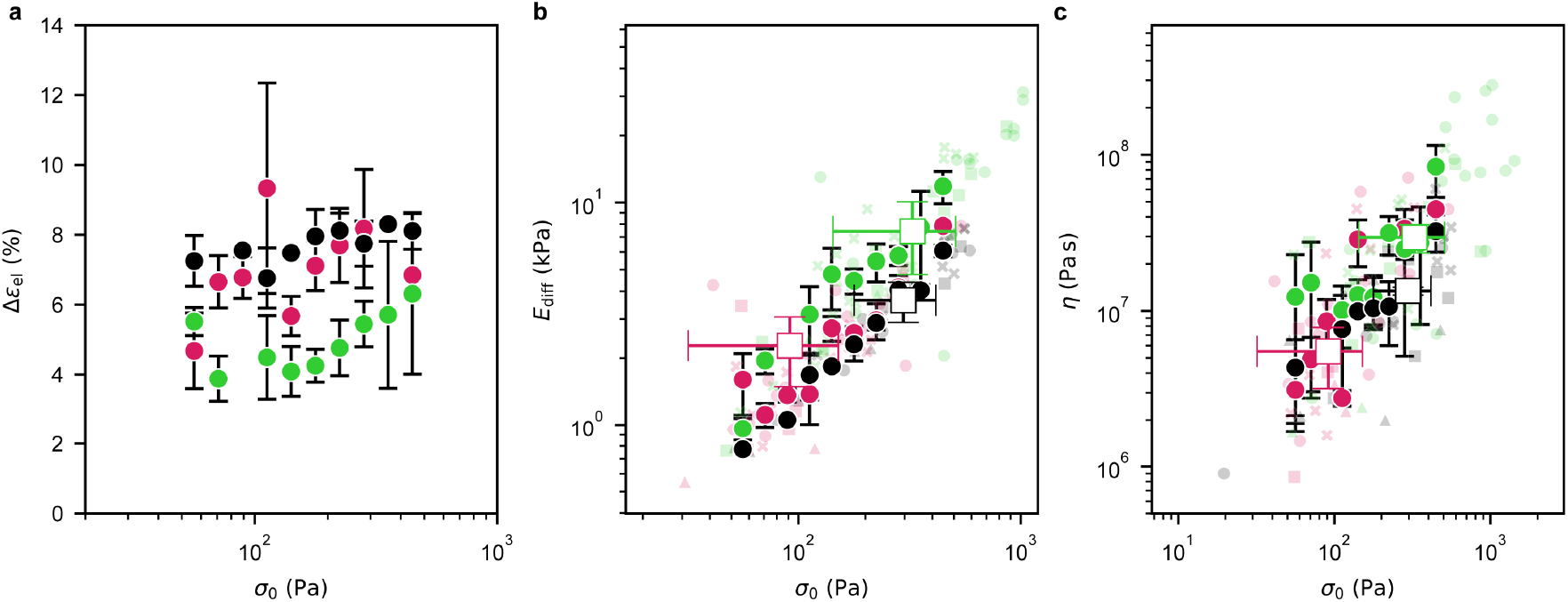
Pel abundance determines inter-strain rheological differences. **a**, Elastic contribution Δ*ε*_el_ to the strain increment for the three strains, binned and averaged within 10 intervals of *σ*_0_ (filled circles), as in Supplementary Fig. 4. The different colours represent the different PA14 mutants: Δpel (magenta), wild-type (black) and ΔwspF (green). The error bars are the standard error of the mean. This plot shows that the strain increment is independent of Δ*σ/σ*_0_. **b**,**c**, Semitransparent markers show the raw data reported in Fig. 2. Fully opaque markers represent values of *E*_diff_ (**b**) and *η* (**c**) binned and averaged as in panel **a**. The colour code is the same as in panel **a**, and the error bars are the standard error of the mean. Data show that, within a given axial stress interval, the Pel-overproducer strain ΔwspF displays significantly higher values of *E*_diff_, while the Pel-deficient Δpel strain and the wild-type have comparable values within the statistical significance of our dataset (**b**). The same trend generally holds for *η* (**c**). The open square markers represent the average values for streamers grown at *U*_gr_ = 2.1 mm*/*s. The error bars are the standard error of the mean. Different edge colours correspond to different mutants, following the same colour code as above. The data show that wild-type and ΔwspF grown at *U*_gr_ = 2.1 mm*/*s are subjected to approximately the same average axial stresses. Thus, the differences in *E*_diff_ and *η* cannot be attributed to the stress hardening behaviour but rather to differences in Pel abundance (Supplementary Fig. 6). In the case of the Δpel mutant, the average axial stress is significantly lower than those of the other strains. This observation suggests that the differences between the average values measured for the wild-type and Δpel mutants result mainly from the stress-hardening behaviour when testing streamers at a fixed *U* rather than at fixed *σ*. In light of these results, we can conclude that there are no significant rheological differences between the Δpel and wild-type matrices at fixed *σ*_0_, in agreement with composition results reported in Supplementary Fig. 6.

**Supplementary Fig. 6.**
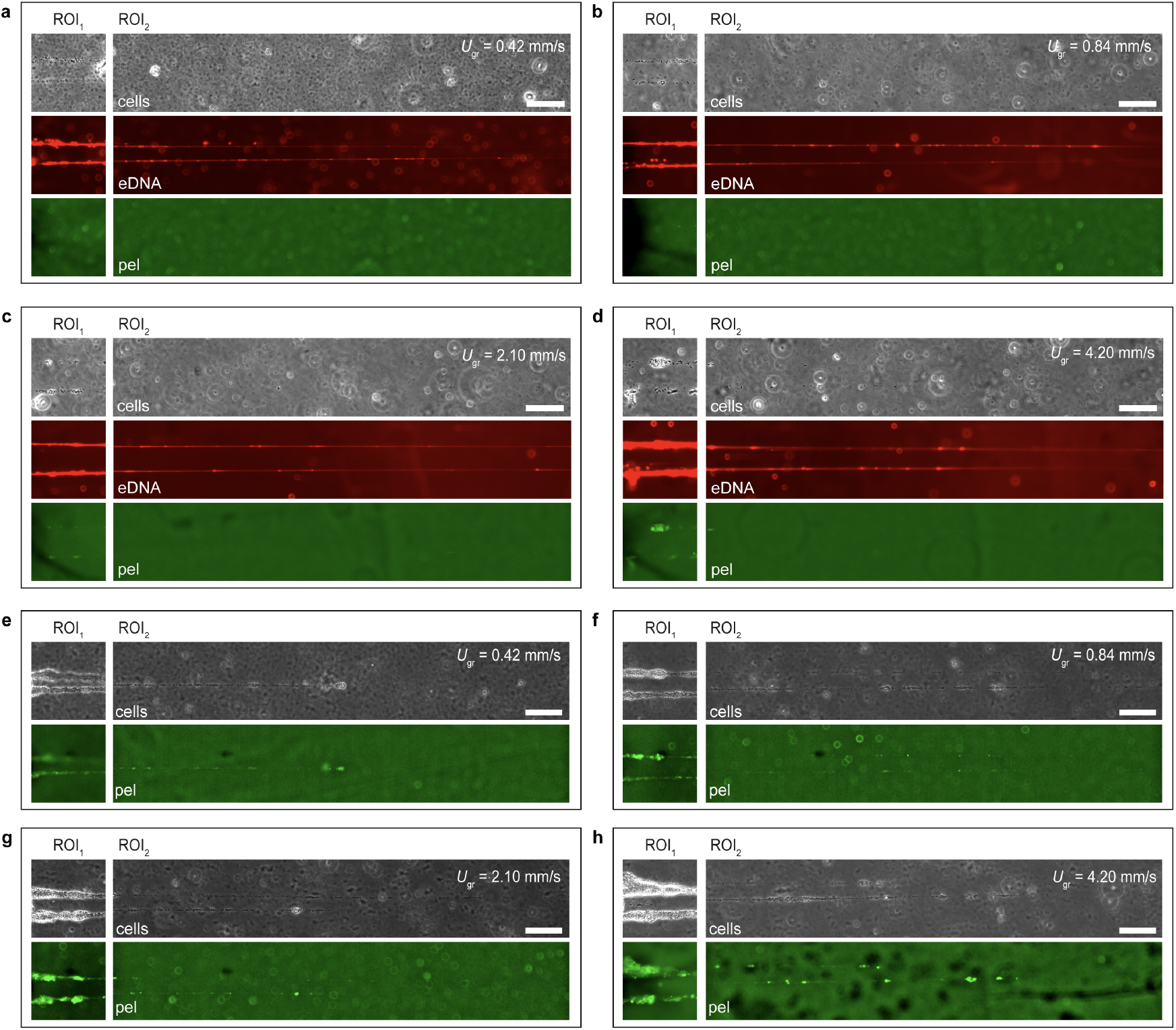
Pel and eDNA distributions along PA14 wild-type and ΔwspF biofilm streamers. a-d, Representative phase contrast and fluorescence images of ROI_1_ and ROI_2_ of PA14 wild-type streamers grown at average flow velocities *U*_gr_ equal to 0.42 mm*/*s (a), 0.84 mm*/*s (b), 2.10 mm*/*s (c) and 4.20 mm*/*s (d). The red fluorescence images show the mCherry signal collected from eDNA stained with PI. eDNA constitutes the backbone of the streamers and is detected both in ROI_1_ and ROI_2_. The green fluorescence images show the GFP signal collected from Pel stained with WFL. Pel is mainly colocalised with the cell aggregates and its abundance in ROI_1_ increases with increasing *U*_gr_. The increase in Pel secretion by cells subjected to higher flow velocities is consistent with the mechanosensing-driven increase in intracellular c-di-GMP levels [14, 15]. Staining with WFL does not detect Pel in ROI_2_ in all flow conditions 5. **e**-**h**, Representative phase contrast and fluorescence images of ROI_1_ and ROI_2_ of PA14 ΔwspF streamers grown at average flow velocities *U*_gr_ equal to 0.42 mm*/*s (e), 0.84 mm*/*s (f), 2.10 mm*/*s (g) and 4.20 mm*/*s (h). The GFP signal shows that Pel is detected both in ROI_1_ and ROI_2_. Pel staining was performed on three independent biological replicates for both PA14 wild-type and ΔwspF with consistent results. Overall, Pel staining results are compatible with the inter-strain rheological differences reported in Supplementary Fig. 5

**Supplementary Fig. 7.**
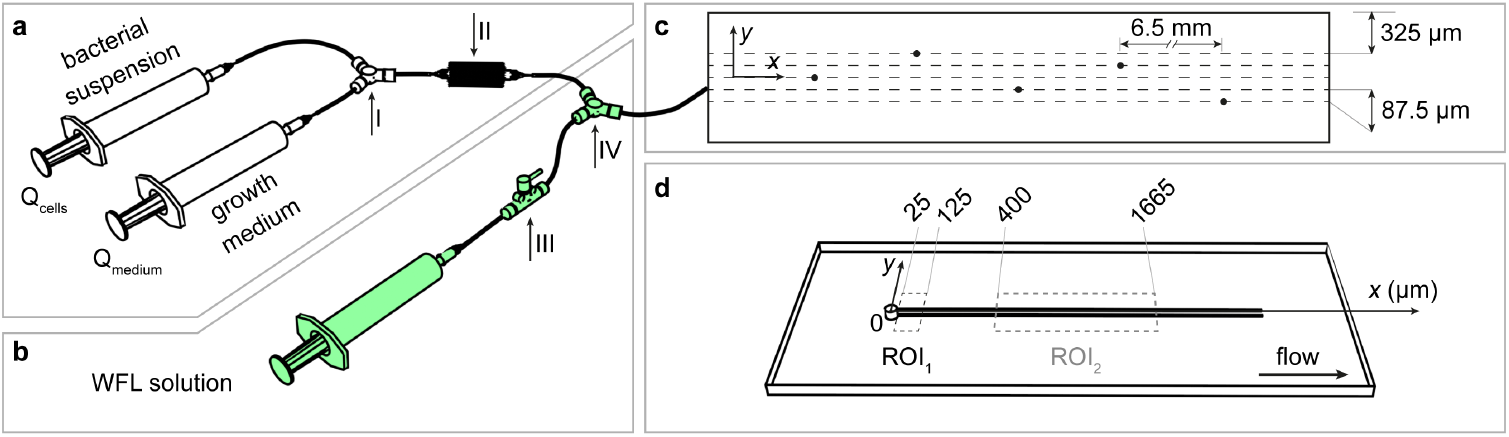
Schematic of the experimental setup. **a**, Setup used to grow biofilm streamers at different *U*_gr_ and constant flux of cells. The flow rate of the bacterial suspension is *Q*_cells_ = 0.06 ml*/*h in all flow conditions. The values of the flow rate *Q*_medium_ of culture medium used in the different flow conditions are reported in Supplementary Table 1. Only the syringe with the bacterial suspension is used in the channel with the lowest flow rate. The total flow rate values *Q*_gr_ = *Q*_cells_ + *Q*_medium_ during streamer growth in the different flow conditions and the corresponding average flow velocities *U*_gr_ are reported in Supplementary Table 1. Arrow I indicates the Y connector used to join the streams from the two syringes. Arrow II indicates the straight connector used to homogenise the sample before the channel inlet. **b**, Additional stream used to flow WFL and stain Pel in the matrix of PA14 wild-type and ΔwspF streamers. This flow line is present only in the WFL staining experiments. Arrow III points at the shut-off valve that allow delayed connection of the syringe at *t* = 15 h, preventing overnight degradation of WFL. After imaging of Pel, the WFL syringe is replaced with a syringe containing a PI solution. Arrow IV points at the Y connector used to join the stream from this additional syringe to the main one. **c**, Top-view schematic of the pillar arrangement inside a channel of the microfluidic platform. We modified the arrangement from our previous works [2, 3] to enhance streamer formation. In this work, pillars are not aligned along the centerline of the channel but evenly spaced along the *y*-axis, with an inter-pillar distance of 87.5 µm. In the current configuration, pillars are evenly spaced along the width of the channel and do not intercept the same flow lines. The distance between the outermost pillars and the channel walls is 325 µm. The distance between pillars in the *x* direction is 6.5 mm (not in scale in the schematic). 3D CFD simulations confirm that the flow field around each pillar is equivalent to that in the original device. **d**, Schematic of two streamers tethered to a micropillar. The dashed rectangles mark ROI_1_ and ROI_2_.

**Supplementary Fig. 8.**
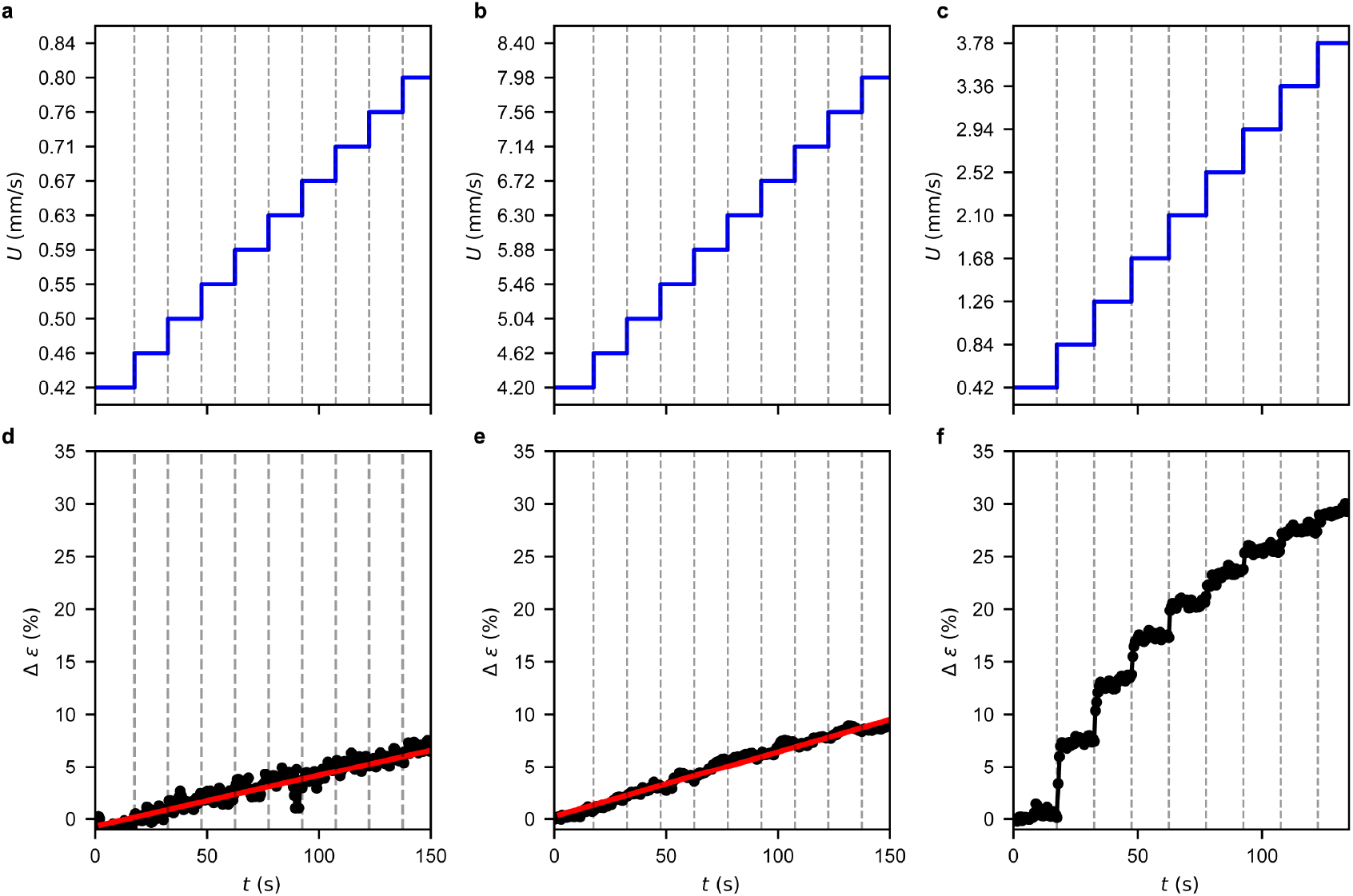
Streamers under prestress *σ*_0_ linearly respond to an axial stress increment Δ*σ* = *σ*_0_. **a, b, d, e**, Tests performed to verify that the response of the streamers subjected to an axial stress increment Δ*σ* = *σ*_0_ is locally linear in a neighbourhood of *σ*_0_. The average velocity *U* in the channel is increased from *U*_gr_ to 2*U*_gr_ by small steps Δ*U* = *U*_gr_*/*10. The time duration of each step is 15 s. Simultaneously, the corresponding strain increment Δ*ε* is measured. Panels **a** and **b** show the average flow velocity inside the microchannel as a function of time during such tests, for the cases *U*_gr_ = 0.42 mm*/*s and *U*_gr_ = 4.20 mm*/*s, respectively. Panel **d** shows the typical deformation curve measured for a streamer grown at *U*_gr_ = 0.42 mm*/*s and subjected to the flow steps in **a**; panel **e** shows the same quantity measured for a streamer grown at *U*_gr_ = 4.20 mm*/*s and subjected to the flow steps in **b**. As shown by the linear fits (red lines), the deformation is linear with time, both for this study’s lowest and highest flow conditions. This implies that the response of the streamers to the applied axial stress is linear between *σ*_0_ and *σ*_0_ + Δ*σ*. **c, f** Experiment showing the nonlinear response of a streamer subjected to a stress increment Δ*σ > σ*_0_. **c** Average velocity *U* in the channel, increasing from *U*_gr_ = 0.42 mm*/*s to 9*U*_gr_ by steps with magnitude Δ*U* = *U*_gr_. The time duration of each step is 15 s. **f** The corresponding steps in strain increment Δ*ε* decrease with increasing *σ*, showing that the response of the streamer to the applied axial stress is nonlinear when Δ*σ > σ*_0_. The decreasing magnitude of the steps of the strain increment Δ*ε* with increasing *σ* clearly demonstrates the stress-hardening behaviour of the streamer.

**Supplementary Fig. 9.**
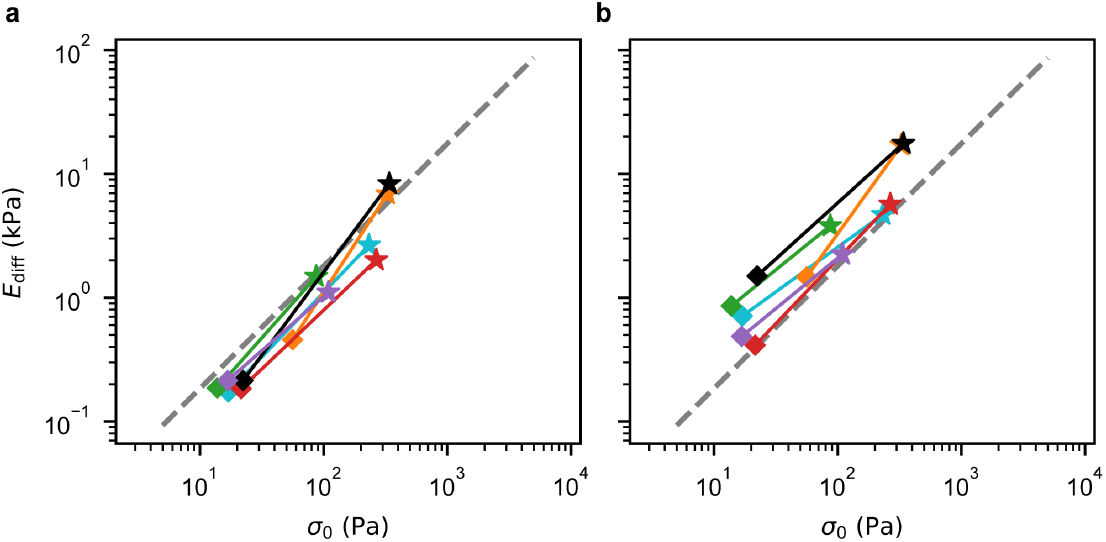
Effects of fluid-structure interaction on the stress hardening behaviour of PA14 wild-type streamers. **a**, Data reported in 3b; each colour identifies two sequential tests related to a same streamer. **b**, Data of panel **a** calculated accounting for the effect of fluid-structure interaction (FSI), following the iterative scheme presented in [2]. The scheme was arrested at the first iteration since higher-order iterations have a negligible effect on the FSI correction, as shown in [2]. The sequential tests in panel **a** have an average slope 1.15 ± 0.10, while the ones in panel **b** 0.95 ± 0.10. The dashed grey lines in panels **a** and **b** show the fit of the data reported in Fig. 3b, with power-law behaviour *σ*^0.99^. These data show that correcting for FSI does not significantly impact the stress-hardening behaviour measured in our experiments.

**Supplementary Fig. 10.**
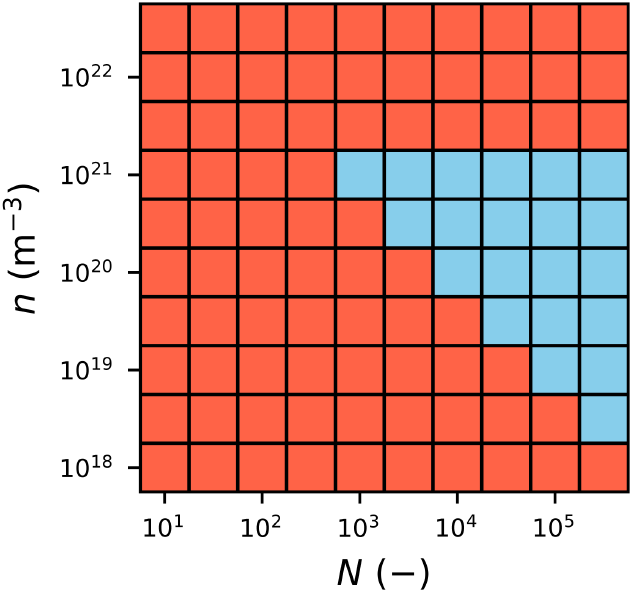
Worm-like chain network model parameter sweep over *N* and *n*. Map showing combinations of *n* and *N* parameter values tested in the worm-like chain network model under uniaxial extension. Each grid cell corresponds to a specific (*N, n*) pair. Blue squares indicate parameter combinations for which the model exhibits linear scaling of the differential Young’s modulus within the prestress range probed experimentally (*σ*_0_ ∈ [4.6 Pa, 4.6 kPa]); red squares indicate combinations for which the model deviates from linearity within that range. Based on this analysis, the parameter region compatible with the observed experimental scaling is defined by *n* ≤10^21^ m^*−*3^ and *N* ≥ 1.5 × 10^24^ m^3^ *n*^*−*1^. These minimal constraints are sufficient for reproducing the experimental scaling without requiring detailed assumptions on the molecular structure of the streamers.

**Supplementary Fig. 11.**
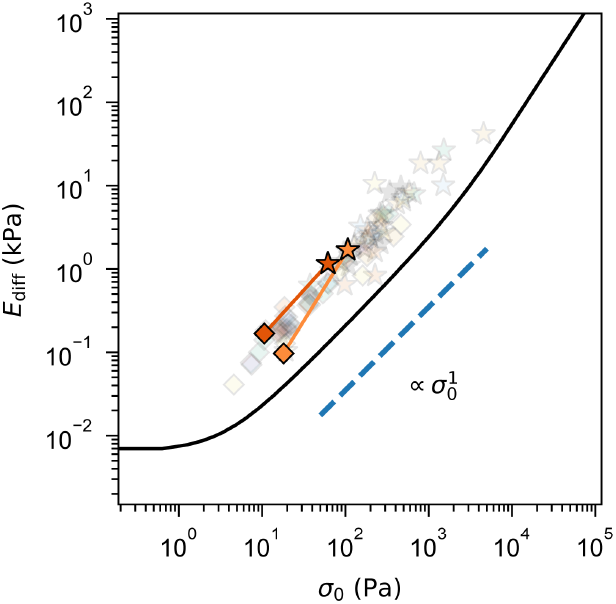
Treatment with RNase buffer at 37 °C does not alter the stress-hardening behaviour of biofilm streamers. Differential Young’s moduli *E*_diff_ measured in sequential mechanical tests after treatment with RNase buffer (50 mM Tris-HCl, pH 7.5), plotted as a function of the axial stress *σ*_0_, (markers in shades of orange). Solid lines connect data points from the same portion of streamer tested under two different prestress conditions *σ*_0_. Tests are conducted on 15-hour-old streamers with initial flow velocities *U*_0_ = *U*_gr_ = 2.10 mm*/*s (stars) and *U*_0_ = *U*_gr_*/*5 = 0.42 mm*/*s (diamonds), corresponding to *σ*_0_ = *σ*(*x*; *U*_gr_)_12_ and *σ*_0_ = *σ*(*x*; *U*_gr_*/*5)_12_, respectively. Before the tests, streamers were treated with proteinase RNase buffer for 1 h. Data show stress-hardening with scaling *E*_diff_ *σ*^1^ (dashed blue line), in agreement with previous results and with the model reported in Fig. 4i (black line), demonstrating that the effect observed in Fig. 5 is due to RNA degradation. For reference, semitransparent markers indicate data from previous experiments on untreated samples (Fig. 4i).

**Supplementary Fig. 12.**
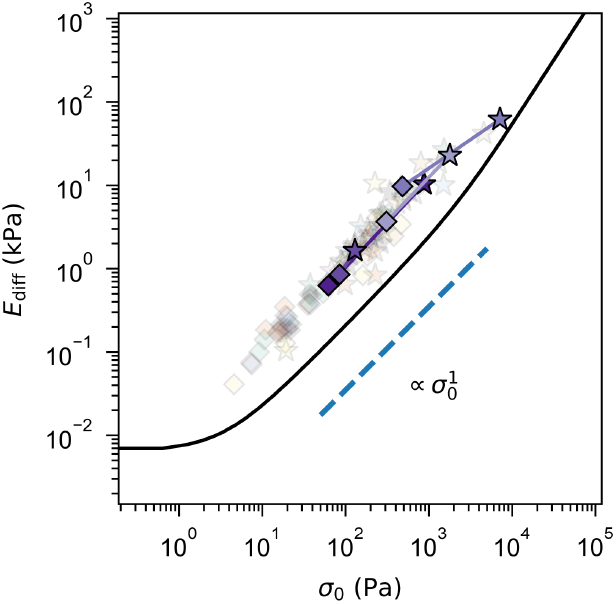
Proteinase K does not affect the stress-hardening behaviour of biofilm streamers. Differential Young’s moduli *E*_diff_, obtained from sequential mechanical tests after proteinase K treatment, plotted as a function of the axial stress *σ*_0_, (markers in shades of purple). Solid lines connect results from the same portion of streamer tested under two prestress conditions *σ*_0_. Tests are conducted on 15-hour-old streamers with initial flow velocities *U*_0_ = *U*_gr_ = 2.10 mm*/*s (stars) and *U*_0_ = *U*_gr_*/*5 = 0.42 mm*/*s (diamonds), corresponding to *σ*_0_ = ⟨*σ*(*x*; *U*_gr_)⟩_12_ and *σ*_0_ = ⟨*σ*(*x*; *U*_gr_*/*5)⟩_12_, respectively. Before the tests, stramers were treated with proteinase K for 1 h. Data show stress-hardening with scaling 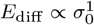 (dashed blue line), in agreement with previous results and with the model reported in Fig. 4i (black line). Semitransparent markers show the data of Fig. 4i, with no proteinase K treatment.

**Supplementary Table 1.** Flow parameters for the four channels of the microfluidic platform, used to simultaneously test streamer growth under different flow conditions. The same bacterial batch is used across all channels to minimise biological variability. *Q*_cells_ is the flow rate of the bacterial suspension, *Q*_medium_ is that of the culture medium, and *Q*_gr_ is the total flow rate inside the channel. The average flow velocity is calculated as *U*_gr_ = *Q*_gr_*/*HW, where H = 40 µm and W = 1000 µm are the channel height and width, respectively. The Reynolds number Re for each condition is calculated as Re = *ρU*_gr_D, where *ρ* is the density of water, and D = 50 µm is the pillar diameter.

| | $Q_{\text{cells}}$ (ml/h) | $Q_{\text{medium}}$ (ml/h) | $Q_{\text{gr}}$ (ml/h) | $U_{\text{gr}}$ (mm/s) | $\text{Re}$ (—) |
| --- | --- | --- | --- | --- | --- |
| channel 1 | 0.06 | 0.00 | 0.06 | 0.42 | 0.02 |
| channel 2 | 0.06 | 0.06 | 0.12 | 0.84 | 0.04 |
| channel 3 | 0.06 | 0.24 | 0.30 | 2.10 | 0.10 |
| channel 4 | 0.06 | 0.54 | 0.60 | 4.20 | 0.20 |

## Acknowledgements

We thank Prof. Leo Eberl, Prof. Jan Vermant, Dr. Sam Charlton and Dr. Steffen Geisel for the insightful discussions; Prof. Leo Eberl for providing the *B. cenocepacia* strain and Prof. Paul D. Fey for the *S. epidermidis* strain; Ela Burmeister for the technical support; and support from the Swiss National Science Foundation PRIMA Grant 179834 (to E.S.) and the Institut Universitaire de France (L.C.). Finally, we acknowledge and honour the memory of Prof. Paul Stoodley, whose pioneering work in streamer formation and rheology has greatly influenced and inspired this research.

## Author contributions

G.S. and E.S. designed the research. G.S. performed the experiments and analysed all the data. G.S. and T.R. contributed to modelling the differential Young’s modulus of streamers. T.R. provided oversight on the analysis of streamer growth. D.T. and L.C. provided oversight on the rheological characterization, data interpretation, and modelling. E.S. provided oversight on data analysis, interpretation, and modelling. G.S. and E.S. wrote the manuscript. All authors edited and commented on the manuscript. E.S. acquired funding.

## Supplementary information

### Elastic stress-hardening behaviour of a DNA network under uniaxial extension

To investigate the origin of the stress-hardening elasticity of the streamers, we compared the power law behaviour observed in our experiments (Fig. 1a, 4i) with the behaviour expected for a 3D network of DNA molecules under uniaxial extension. We describe the force-extension curve of single DNA molecules constituting the network with the following worm-like chain interpolated formula [1], which takes into account the entropic contribution to DNA elasticity:

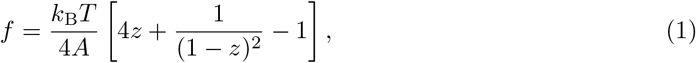

where *z* is the extension ratio of the molecule, *A* is the persistence length of DNA, *k*_B_ is the Boltzmann’s constant and *T* is the temperature. The extension ratio is defined as *z* = *r/L*, where *r* is the end-to-end distance of the DNA molecule and *L*_*c*_ its contour length. The free energy *W* of the DNA molecule subjected to such a force can be calculated by integrating equation 1:

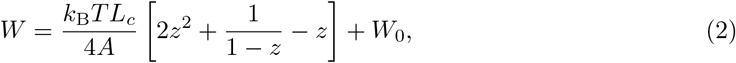

where *W*_0_ is a constant [2]. This single-molecule free energy can be used to find the three-dimensional free energy of a network of chains, employing the eight-chain model developed in [3]. This model assumes that a microscopic cube element of a network with edge *a*_0_ has a cross-link in the centre and eight chains extending from the cross-link to the eight corners of the cube. This description assumes that the network is isotropic in the unstrained state, and is equivalent to averaging the contribution of a single chain over eight spatial orientations. This allows finding a relation between the end-to-end distance *r* of the chains within the network and the invariants of the Cauchy strain tensor, describing the macroscopic state of deformation of the network. In the unstrained state, the end-to-end distance of each chain is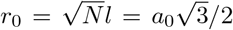, where *N* is the number of freely jointed segments of length *l* in a chain between neighbouring cross-links. For an arbitrary strained state where the edges of the microscopic cube are *a*_*i*_ = *a*_0_ *λ*_*i*_ (*i* = *x, y, z*), the *r* of each chain is:

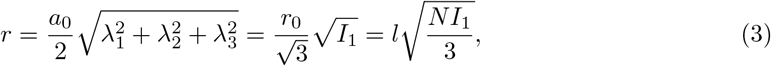

where 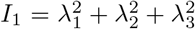 is the first invariant of the Cauchy strain tensor and *λ*_1_, *λ*_2_, *λ*_3_ are the principal stretches. By substituting eq. 3 into 2, the three-dimensional free energy of a molecule within the network takes the form:

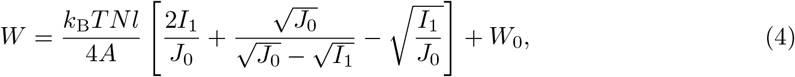

where *J*_0_ = 3*N* [2]. This energy is the contribution of a single worm-like chain within the network. Thus, the energy of the whole network is obtained by mutiplying eq. 4 by the number of chains per unit volume *n*:

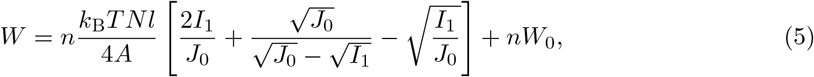

In the case of uniaxial extension, assuming network incompressibility, the principal stretches are:

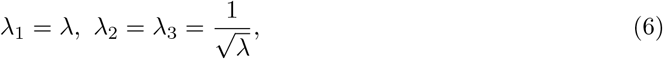

where *λ* is the stretch ratio in the direction of extension. Therefore, the first invariant is:

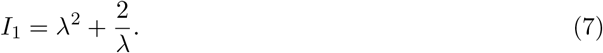

In this case, the true stress (calculated with respect to the deformed cross section) in the direction of extension is [2]:

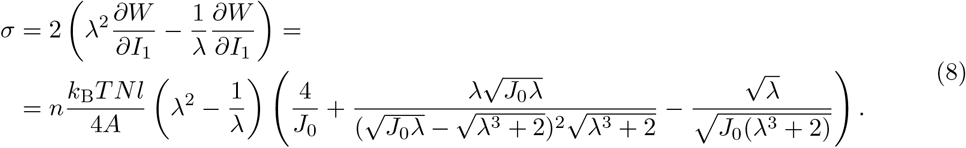

Due to the finite chain extensibility, the stress diverges when *λ → λ*^*\**^, where *λ*^*\**^ is the stretch ratio such that

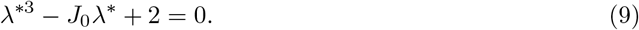

From equation 8, the differential Young’s modulus *E*_diff_ of the network can be obtained as [4]:

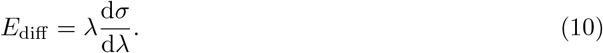

From equations 8 and 10 it is then possible to obtain the functional relation between *E*_diff_ and *σ* for a network under uniaxial extension (Fig. 4i). This shows a plateau for small values of *σ*, corresponding to the linear elastic response of the network, and region at large *σ* where *E*_diff_ *∝ σ*^1.5^. When the network is such that the number *N* of freely jointed segments between neighbouring cross-links is high enough (Supplementary Fig. 10), the model predicts a cross-over regime for intermediate values of *σ* where *E*_diff_ *∝ σ*, as observed for the streamers.

For a network of DNA molecules (*A ≈* 50 nm [1]) at room temperature (*T* = 293 K), the cross-over regime coincides with the prestress range spanned in the sequential tests (*σ*_0_ ∈ [4.6 Pa, 4.6 kPa], Fig. 4i) when considering *N* = 1000 and chain density *n* = 1 *×* 10^21^ m^*−*3^. The corresponding contour length between neighbouring cross-links is thus *L*_*c*_ = *Nl* = 50 µm, with the length *l* of a single freely jointed segment chosen as *l* = *A* = 50 nm, satisfying the condition *l* ≤*A* ≤*L*_*c*_ reported in [2]. Supplementary Fig. 10 reports the more general conditions under which the cross-over regime contains and is larger than the prestress range spanned in the sequential tests (Fig. 4i): *n ≤* 10^21^ m^*−*3^ and *N ≥* 1.5 *×* 10^24^ m^3^ *· n*^*−*1^

## Notes

### Competing Interest Statement

The authors have declared no competing interest.

### Summary of Updates

We have incorporated two specific discussions: 1.Clarification on Pel polysaccharides: We expanded the discussion to clarify the consistency of our current findings with our earlier work (Secchi et al., PNAS 2022; Savorana et al., Soft Matter 2022). We emphasize that while Pel concentration modulates the magnitude of viscoelastic properties at a given prestress, it does not determine the presence or functional dependence of stress-hardening. 2.Nature of eDNA cross-linking: We added a discussion of possible cross-linking mechanisms, in particular the potential role of Holliday junctions between eDNA molecules, as suggested in recent literature. We highlight that the reported cross-link length scale is consistent with that emerging from our modelling.

