## Supplementary information for "Stress-hardening behaviour of biofilm streamers"

### Elastic stress-hardening behaviour of a DNA network under uniaxial extension

To investigate the origin of the stress-hardening elasticity of the streamers, we compared the power law behaviour observed in our experiments (Fig. 1a, 4i) with the behaviour expected for a 3D network of DNA molecules under uniaxial extension. We describe the force-extension curve of single DNA molecules constituting the network with the following worm-like chain interpolated formula [1], which takes into account the entropic contribution to DNA elasticity:

$$f = \frac{k_\mathrm{B}T}{4A}\left[4z + \frac{1}{(1-z)^2} - 1\right], \tag{1}$$

where $z$ is the extension ratio of the molecule, $A$ is the persistence length of DNA , $k_\mathrm{B}$ is the Boltzmann's constant and $T$ is the temperature. The extension ratio is defined as $z = r/L$, where $r$ is the end-to-end distance of the DNA molecule and $L_c$ its contour length. The free energy $W$ of the DNA molecule subjected to such a force can be calculated by integrating equation 1:

$$W = \frac{k_\mathrm{B}TL_c}{4A}\left[2z^2 + \frac{1}{1-z} - z\right] + W_0, \tag{2}$$

where $W_0$ is a constant [2]. This single-molecule free energy can be used to find the three-dimensional free energy of a network of chains, employing the eight-chain model developed in [3]. This model assumes that a microscopic cube element of a network with edge $a_0$ has a cross-link in the centre and eight chains extending from the cross-link to the eight corners of the cube. This description assumes that the network is isotropic in the unstrained state, and is equivalent to averaging the contribution of a single chain over eight spatial orientations. This allows finding a relation between the end-to-end distance $r$ of the chains within the network and the invariants of the Cauchy strain tensor, describing the macroscopic state of deformation of the network. In the unstrained state, the end-to-end distance of each chain is $r_0 = \sqrt{N}l = a_0\sqrt{3}/2$, where $N$ is the number of freely jointed segments of length $l$ in a chain between neighbouring cross-links. For an arbitrary strained state where the edges of the microscopic cube are $a_i = a_0\,\lambda_i$ $(i = x, y, z)$, the $r$ of each chain is:

$$r = \frac{a_0}{2}\sqrt{\lambda_1^2 + \lambda_2^2 + \lambda_3^2} = \frac{r_0}{\sqrt{3}}\sqrt{I_1} = l\sqrt{\frac{NI_1}{3}}, \tag{3}$$

where $I_1 = \lambda_1^2 + \lambda_2^2 + \lambda_3^2$ is the first invariant of the Cauchy strain tensor and $\lambda_1$, $\lambda_2$ , $\lambda_3$ are the principal stretches. By substituting eq. 3 into 2, the three-dimensional free energy of a molecule within the network takes the form:

$$W = \frac{k_\mathrm{B}TNl}{4A}\left[\frac{2I_1}{J_0} + \frac{\sqrt{J_0}}{\sqrt{J_0} - \sqrt{I_1}} - \sqrt{\frac{I_1}{J_0}}\right] + W_0, \tag{4}$$

where $J_0 = 3N$ [2]. This energy is the contribution of a single worm-like chain within the network. Thus, the energy of the whole network is obtained by mutiplying eq. 4 by the number of chains per unit volume $n$:

$$W = n\frac{k_{\mathrm{B}}TNl}{4A}\left[\frac{2I_1}{J_0} + \frac{\sqrt{J_0}}{\sqrt{J_0}-\sqrt{I_1}} - \sqrt{\frac{I_1}{J_0}}\right] + nW_0, \tag{5}$$

In the case of uniaxial extension, assuming network incompressibility, the principal stretches are:

$$\lambda_1 = \lambda,\ \ \lambda_2 = \lambda_3 = \frac{1}{\sqrt{\lambda}}, \tag{6}$$

where $\lambda$ is the stretch ratio in the direction of extension. Therefore, the first invariant is:

$$I_1 = \lambda^2 + \frac{2}{\lambda}. \tag{7}$$

In this case, the true stress (calculated with respect to the deformed cross section) in the direction of extension is [2]:

$$\begin{aligned}\sigma &= 2\left(\lambda^2\frac{\partial W}{\partial I_1} - \frac{1}{\lambda}\frac{\partial W}{\partial I_1}\right) = \\ &= n\frac{k_{\mathrm{B}}TNl}{4A}\left(\lambda^2 - \frac{1}{\lambda}\right)\left(\frac{4}{J_0} + \frac{\lambda\sqrt{J_0\lambda}}{(\sqrt{J_0\lambda}-\sqrt{\lambda^3+2})^2\sqrt{\lambda^3+2}} - \frac{\sqrt{\lambda}}{\sqrt{J_0(\lambda^3+2)}}\right).\end{aligned} \tag{8}$$

Due to the finite chain extensibility, the stress diverges when $\lambda \to \lambda^*$, where $\lambda^*$ is the stretch ratio such that

$$\lambda^{*3} - J_0\lambda^* + 2 = 0. \tag{9}$$

From equation 8, the differential Young's modulus $E_{\mathrm{diff}}$ of the network can be obtained as [4]:

$$E_{\mathrm{diff}} = \lambda\frac{\mathrm{d}\sigma}{\mathrm{d}\lambda}. \tag{10}$$

From equations 8 and 10 it is then possible to obtain the functional relation between $E_{\mathrm{diff}}$ and $\sigma$ for a network under uniaxial extension (Fig. 4i). This shows a plateau for small values of $\sigma$, corresponding to the linear elastic response of the network, and region at large $\sigma$ where $E_{\mathrm{diff}} \propto \sigma^{1.5}$. When the network is such that the number $N$ of freely jointed segments between neighbouring cross-links is high enough (Supplementary Fig. 10), the model predicts a cross-over regime for intermediate values of $\sigma$ where $E_{\mathrm{diff}} \propto \sigma$, as observed for the streamers.

For a network of DNA molecules ($A \approx 50\,\mathrm{nm}$ [1]) at room temperature ($T = 293\,\mathrm{K}$), the cross-over regime coincides with the prestress range spanned in the sequential tests ($\sigma_0 \in [4.6\,\mathrm{Pa}, 4.6\,\mathrm{kPa}]$, Fig. 4i) when considering $N = 1000$ and chain density $n = 1 \times 10^{21}\,\mathrm{m}^{-3}$. The corresponding contour length between neighbouring cross-links is thus $L_c = Nl = 50\,\mu\mathrm{m}$, with the length $l$ of a single freely jointed segment chosen as $l = A = 50\,\mathrm{nm}$, satisfying the condition $l \leq A \leq L_c$ reported in [2]. Supplementary Fig. 10 reports the more general conditions under which the cross-over regime contains and is larger than the prestress range spanned in the sequential tests (Fig. 4i): $n \leq 10^{21}\,\mathrm{m}^{-3}$ and $N \geq 1.5 \times 10^{24}\,\mathrm{m}^3 \cdot n^{-1}$
